# A closely related pair of superoxide dismutase isozymes from *Staphylococcus aureus* show distinct stabilities and proton-exchange dynamics

**DOI:** 10.1101/2025.10.27.684957

**Authors:** Mariam Esmaeeli, Lorna Nikolić, Rafał Mazgaj, Swati Das, Lilia Zhukova, Michał Dadlez, Kevin J. Waldron

**Affiliations:** Institute of Biochemistry & Biophysics, Polish Academy of Science, Warsaw 02-106, Poland; Doctoral School of Molecular Biology and Biological Chemistry, Institute of Biochemistry and Biophysics, Polish Academy of Science, Warsaw 02-106, Poland

**Keywords:** Superoxide dismutase, protein stability, hydrogen-deuterium exchange, circular dichroism

## Abstract

Changes in biochemical properties, caused by iterative mutations in amino acid sequence, underlie the alterations in protein function over time that underpin the evolutionary process. An example is the switching of an enzyme’s reliance from one essential metal to an alternative as their catalytic cofactor. We previously described such a neofunctionalisation in *Staphylococcus aureus*, which altered a superoxide dismutase (SOD) enzyme from being an ancestral manganese-dependent (MnSOD) into an extant isozyme that can equally utilise either manganese or iron, termed cambialism (camSOD). Yet it’s unclear whether camSOD emergence involved selection solely for cofactor flexibility or whether other biochemical properties also diverged during neofunctionalisation. Here, we have investigated an independent biochemical property of the *S. aureus* SODs, their structural stability. We demonstrate that the neofunctionalised camSOD exhibits increased stability relative to the ancestral MnSOD. *S. aureus* camSOD is more resistant to both chemical and thermal unfolding *in vitro*. Crucially, while both isozymes possess a stable ‘core’ at the heart of their fold, consisting of regions of the protein localised around the metal cofactor that resist hydrogen-deuterium exchange when exposed to isotopically labelled solvent, this core is larger and more exchange-resistant in camSOD than MnSOD. Thus, during the recent divergence of this SOD pair, two distinct biochemical properties have undergone substantial and rapid evolutionary change. This study paves the way for investigations of the structural and functional relationship between these properties, a SODs metal-preference and stability, and of how these properties were concomitantly selected during neofunctionalisation in the *S. aureus* lineage.

## Introduction

Protein adaptation is a crucial aspect of evolution, whereby iterative changes in the sequence of a polypeptide give rise to altered biochemical properties. The sequence changes are presumed to be primarily the result of random mutations that occur in the DNA sequence during replication, but their selection is non-random. Mutations that have deleterious effects, for example disrupting the protein’s biochemical function or stability, are expected to be negatively selected and driven out of the gene pool, whereas those with beneficial effects will be positively selected in the population. Notably, the selection of novel proteins can be multifactorial, with a new sequence being selected on the basis of one or more beneficial traits. Such evolutionary changes could optimise multiple distinct biochemical properties (for example, the reaction catalysed and the protein’s stability) simultaneously, serially, or even change one property at the expense of another where the latter property is under less stringent selection.

An example of the evolutionary adaptation of a biochemical property of proteins is seen in the switching of the preferred metal cofactor that we have observed in members of the iron- or manganese-dependent superoxide dismutase (SOD) family (SodFM) (1). The SodFMs are an important cellular defence against the reactive oxygen species, superoxide, that are widely distributed across genomes. Their catalytic metal preference for either iron, manganese, or for either metal ion (termed cambialism), is a key biochemical parameter that defines their properties *in vitro* and *in vivo* (2). Nonetheless, the structures of all characterised SodFMs are remarkably similar (1, 3–6), regardless of their metal-preference. Only small changes to the primary sequence, altering the chemistry of residues that lie close to the catalytic metal ion within the final folded structure, are needed to change the catalytic metal-preference of SodFMs (1, 7–11). These changes are found to make no significant alterations to the overall structural fold of the enzymes (7–9, 11), perhaps because changes that do disrupt the fold are negatively selected. These minor changes within the metal’s secondary coordination sphere are proposed to alter the enzyme’s catalytic metal-preference through modulation of its Lewis acidity or its reduction potential, with the latter process termed redox tuning (12–20). On the other hand, it is not known whether other properties of SodFMs, such as their stability or their interactions with other protein partners, might also have been selected evolutionarily in some systems.

The pair of SodFMs from the Gram-positive bacterium *Staphylococcus aureus* are one of the best characterised examples of protein neofunctionalisation (1, 5, 11, 21, 22). The pair emerged after a gene duplication event in the ancestor of *S. aureus* and its closest relatives, *S. argenteus* and *S. schweitzeri* (11). The ancestral gene, *sodA*, encodes a highly manganese-preferring SodFM isozyme (MnSOD), which is common to all staphylococcal genomes, whereas the duplicate gene, *sodM*, underwent neofunctionalisation (11). This process resulted in the selection of a cambialistic isozyme (camSOD), equally capable of catalysis with manganese or iron (21), which we have shown was achieved by mutations to residues that are located within the metal ion’s secondary coordination sphere (11). The result is that extant *S. aureus* possesses a pair of SodFM paralogues with divergent catalytic metal-preferences but near-identical crystal structures (11, 21). The selection pressure that drove this neofunctionalisation is likely the manganese starvation that pathogenic *S. aureus* experiences when it infects a host (21), due to the manganese-chelating action of the immune protein complex calprotectin (23–25). The change of metal-preference of the camSOD, SodM, was accompanied by a change in the regulation of *sodM* with respect to its *sodA* ancestor (21, 26). However, whether the neofunctionalisation process that resulted in the emergence of camSOD involved selection of just a single protein trait, that of its catalytic metal-preference, or multiple traits is not known.

Extant camSOD has acquired 49 mutations in its protein sequence in comparison to MnSOD (out of 199 amino acids: 24.6% of residues differ) during neofunctionalisation, yet only one or two mutations are needed to substantially change its metal-preference (11). The large number of mutations selected in SodM indicates that other properties of camSOD may also have been under positive selection during its neofunctionalisation. Given the rapid decay of stability that is observed as mutations are combined (27) and the subsequent identification of protein stability as the prime factor that determines the rate of protein evolution (28), we hypothesised that SOD stability might also be under positive selection. During earlier attempts to produce isozymes exclusively loaded with manganese through unfolding refolding protocols (11, 21), we observed that MnSOD and camSOD differed in their unfolding in urea and guanidine. In this study, we have investigated the structural stability of these two closely related SodFMs using a range of biochemical and biophysical assays. Our data demonstrate that the *S. aureus* camSOD, SodM, gained additional structural stability relative to its MnSOD ancestor during the neofunctionalisation process that imparted its cambialistic activity. We discuss the potential implications for the evolutionary adaptation of SodM during the emergence of *S. aureus* from its non-pathogenic ancestors.

## Results

### Unfolding of the S. aureus SodFM enzymes in the presence of chaotropic reagents

We first used tryptophan (Trp) fluorescence emission spectroscopy to interrogate an apparent difference in the unfolding of the *S. aureus* SODs in the presence of chaotropic reagents. The wavelength and intensity of Trp fluorescence emission is known to depend on the environment of the aromatic sidechain (29). The staphylococcal SODs contain 7 Trp residues, and thus we hypothesised that monitoring fluorescence would be a sensitive way to monitor unfolding of the hydrophobic core of SodFMs.

Samples of SodA and SodM, loaded with either Mn or Fe as verified by inductively coupled plasma optical emission spectrometry (ICP-OES), were incubated in either 8 M urea or 6 M guanidinium hydrochloride solutions, or in buffer alone (Fig. 1). As expected, all four untreated forms exhibited emission spectra centred on an emission maximum (λ_max_) at 330-335 nm, consistent with their Trp residues being localised inside the folded protein and protected from exposure to solvent (30). Incubation of MnSOD, loaded with either metal, with either of these chaotropic agents resulted in a red-shift in λ_max_ to the 350-360 nm range, concomitant with an increase in emission intensity (Fig. 1 A,C). These changes in fluorescence emission imply a significant change of environment of the Trp residues of MnSOD resulting in their exposure to solvent, indicating that the hydrophobic core structure of SodA is disrupted by high concentrations of either urea or guanidine. Notably though, the fluorescence changes indicated that either chaotropic agent could efficiently unfold Mn-SodA (Fig. 1A), whereas urea was markedly less effective at unfolding Fe-SodA (Fig. 1C).

**Fig. 1:**
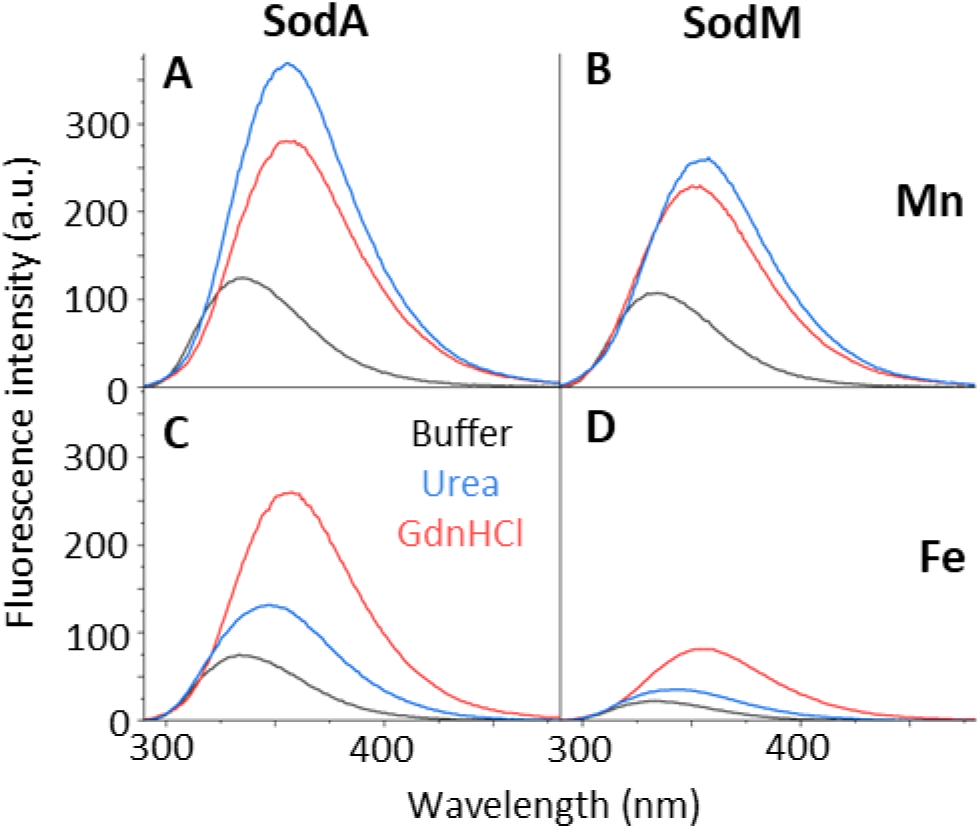
Fluorescence emission of Trp residues in *S. aureus* SODs shows their differential unfolding in urea or guanidine. Trp fluorescence emission was assessed from samples (10 µM) of recombinant (A,C) SodA or (B,D) SodM from *S. aureus*, loaded either with (A,B) manganese or (C,D) iron. By ICP-OES analysis, the Mn-SodA sample used (A) contained 0.860 mole equivalent Mn, 0.002 Fe, 0.002 Zn, and the Fe-SodA sample (B) contained 0.849 mole equivalent Fe, 0.011 Mn, 0.002 Zn, whereas the Mn-SodM sample (C) contained 0.835 Mn, 0.055 Fe, 0.002 Zn, and the Fe-SodM sample (D) contained 1.090 Fe, 0.005 Mn, 0.000 Zn. Triplicate samples were incubated at room temperature for 24 h in either buffer (20 mM Tris, 5 mM EDTA, pH 7.5, 150 mM NaCl - black), in 8 M urea (blue) or in 6 M guanidine prior to spectra being acquired. Measurements were performed at 600 V in 10 mm path length quartz cuvettes, with excitation and emission slits at 5 nm. Triplicate measurements were obtained, subtracting spectra obtained from their respective blank solutions, and averaged.

The fluorescence emission of the camSOD, SodM, showed distinct behaviour in the presence of the chaotropic agents. Whereas Mn-SodM showed both a shift in λ_max_ (358 nm) and intensity after incubation with either urea or guanidine (Fig. 1 B), Fe-SodM showed a much smaller shift in λ_max_ and only a small increase in emission intensity in guanidine, and negligible changes at all in the presence of urea (Fig. 1 D). This indicated that the iron-loaded form of camSOD was highly resistant to unfolding of the hydrophobic core. The intermediate shift of λ_max_ and increase in intensity observed in the presence of guanidine likely reflect a partially unfolded state of Fe-SodM, in which its multiple Trp residues experience differences in solvent exposure, whereas those in SodA are in a more uniform unfolded state exposed to solvent. Collectively, these data indicated that camSOD was more resistant to unfolding in urea than MnSOD in both metal-bound form, but that the Fe-loaded form of camSOD in particular was completely resistant to urea.

Next, we employed circular dichroism (CD) spectroscopy to further explore the unfolding of MnSOD and camSOD in the presence of the chaotropic agents. We followed the change of the CD signal at 222 nm to assess the α-helical content of the proteins after incubation with varying concentrations of urea and guanidine (Fig. 2,3 and Supp. Figs 1-4). The effect of these agents on the proteins and their residual activity were also assessed from SOD activity- and Coomassie blue-stained native gels (Fig. 2,3 and Supp. Figs. 3,4,6).

**Fig. 2:**
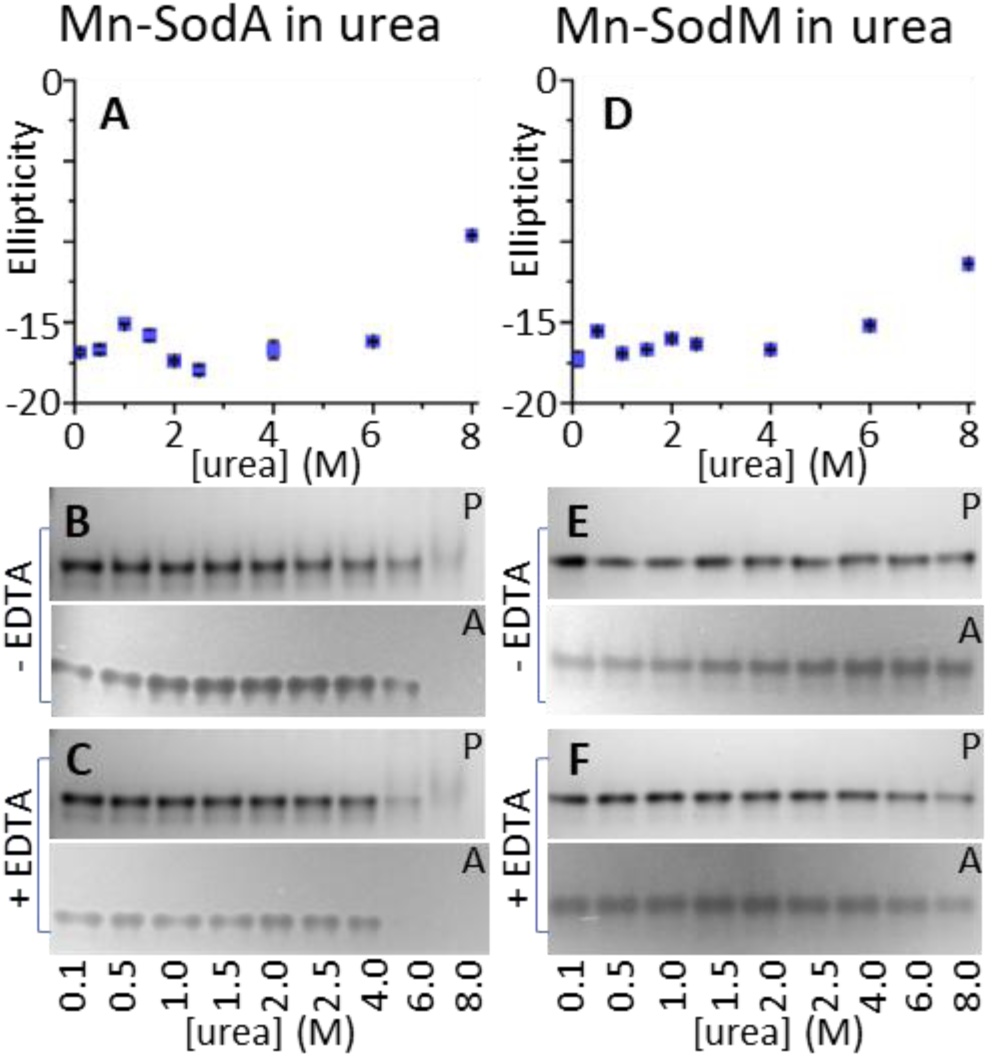
CD spectroscopy shows differential unfolding of manganese-loaded *S. aureus* SODs in urea. Unfolding of the manganese-loaded isoforms of (A-C) SodA or (D-F) SodM by the chaotropic agent urea was assessed by (A,D) measuring their CD spectra and by assessing (B,E) protein (P) stability through Coomassie staining and (C,F) enzymatic activity (A) through NBT/riboflavin staining of native PAGE gels, either before or after incubation of the samples with EDTA. Protein (10 μM) samples were in 50 mM potassium phosphate buffer, pH 7.5. Samples of each protein (Mn-SodA was determined to contain 0.983 mole equivalent Mn, 0.020 Fe, 0.001 Zn; Mn-SodM contained 1.070 mole equivalent Mn, 0.050 Fe, 0.00 Zn) were incubated overnight at 4 °C with different concentrations of urea (0.1 to 8 M) and then their CD spectra were over the 205 to 260 nm range (Supp. Fig. 1). Each sample was measured in technical triplicate and results presented as average molar ellipticity (deg.cm^2^.dmol^-1^) ± standard deviation. EDTA (5 μL of 50 mM) was added to aliquots (2.5 μL) of each urea-incubated sample, incubated for 2 h, then aliquots of both control and EDTA-treated urea-incubated protein samples were resolved on 10 or 12% acrylamide native PAGE, and gels stained with either Coomassie Brilliant Blue for detecting protein or with NBT/riboflavin stain to detect SOD activity. For Coomassie staining, aliquots containing 280 ng protein were loaded of all samples, whereas for activity staining, aliquots containing 17.5 ng of Mn-SodA or 33 ng Mn-SodM were loaded. The uncropped gel images used in this composite figure are shown in Supp. Fig. 6.

**Fig. 3:**
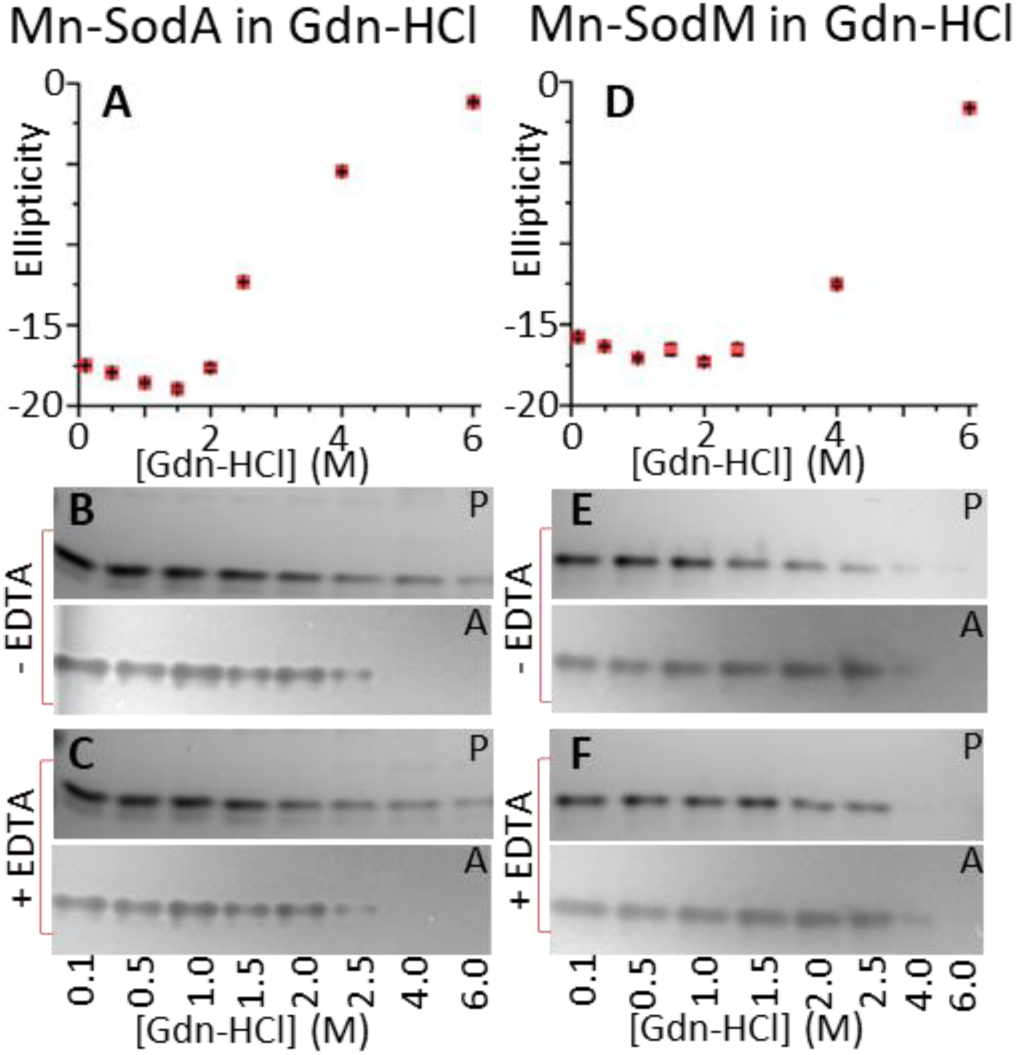
CD spectroscopy shows differential unfolding of manganese-loaded *S. aureus* SODs in guanidine. Unfolding of the manganese-loaded isoforms of (A-C) SodA or (D-F) SodM by the chaotropic agent guanidine was assessed by (A,D) measuring their CD spectra and by assessing (B,E) protein (P) stability through Coomassie staining and (C,F) enzymatic activity (A) through NBT/riboflavin staining of native PAGE gels, either before or after incubation of the samples with EDTA. Protein (10 μM) samples were in 50 mM potassium phosphate buffer, pH 7.5. Samples of each protein (metal content as in Fig. 1) were incubated overnight at 4 °C with different concentrations of guanidine (0.1 to 6 M) and then their CD spectra were over the 205 to 260 nm range (Supp. Fig. 2). Each sample was measured in technical triplicate and results presented as the average molar ellipticity (deg.cm^2^.dmol^-1^) ± standard deviation. EDTA (5 μL of 50 mM) was added to aliquots (2.5 μL) of each guanidine-incubated sample, incubated for 2 h, then aliquots of both control and EDTA-treated guanidine-incubated protein samples were resolved on 10 or 12% acrylamide native PAGE, and gels stained with either Coomassie Brilliant Blue for detecting protein or with NBT/riboflavin stain to detect SOD activity. For Coomassie staining, aliquots containing 280 ng protein were loaded of all samples, whereas for activity staining, aliquots containing 17.5 ng of Mn-SodA or 33 ng Mn-SodM were loaded. The uncropped gel images used in this composite figure are shown in Supp. Fig. 7.

Mn-SodA retained its helical secondary structure up to 6 M urea but showed substantial unfolding in 8 M urea (Fig. 2 A). Notably, the protein band for Mn-SodA and its activity decreased in intensity at 6 M urea, and only a faint band remained visible at 8 M urea (Fig. 2 B). When these samples were exposed to the divalent metal chelator EDTA (50 mM), the protein band intensity decreased, and its activity disappeared completely even at 6 M urea (Fig. 2 C). This indicated that the Mn-loaded MnSOD structure is significantly destabilised by 6 M urea, despite limited loss of helical content, such that the manganese ion bound in the enzyme’s active site can be extracted by EDTA in the presence of the chaotropic agents, but that in the absence of the chelator the structure can refold around the metal ion on entering the gel. On the other hand, Mn-SodM showed less loss of secondary structure in 8 M urea (Fig. 2 D), and these conditions only resulted in a minor loss of activity (Fig. 2 E), even after incubation with EDTA (Fig. 2 F). As was seen by Trp fluorescence (Fig. 1), this indicated that Mn-SodA was more susceptible to unfolding in urea than Mn-SodM. Interestingly, this trend was also observed, albeit not as strongly, when we compared CD spectra of this pair of SODs when loaded with iron (Supp. Fig. 3). CD spectra indicated only small decreases in the α-helical content of both Fe-SodA and Fe-SodM in the presence of urea (Supp. Fig. 3 A, 3 D), and no decrease was seen in the protein or activity of either of these isoforms (Supp. Fig. 3 B,E), even in the presence of EDTA (Supp. Fig. 3 C,F).

The effect of guanidine on α-helical structure was more drastic. In both manganese- (Fig. 3) and iron-loaded forms (Supp. Fig. 4), both SODs showed total loss of helical structure by CD spectroscopy in the presence of 6 M guanidine. However, the effects of lower concentrations of guanidine demonstrated clear differences between the MnSOD and the camSOD. For example, Mn-SodA exhibited decreased helical content at 2.5 M guanidine and near-total loss at 4 M (Fig. 3 A). Consistent with this, activity was decreased at 2.5 M and lost completely at 4 M guanidine, both in the absence (Fig. 3 B) and presence of EDTA (Fig. 3 C). Conversely, Mn-SodM showed no loss of helical content at 2.5 M and only partial loss at 4 M guanidine (Fig. 3 D), which was also reflected in activity (Fig. 3 E,F). A similar trend was observed in the CD spectra of the Fe-loaded forms, with Fe-SodA showing loss of helical content and activity at 4 M guanidine (Supp. Fig. 4 A-C), whereas Fe-SodM showed negligible loss of helical content or activity under the same conditions (Supp. Fig. 4 D-F).

Both spectroscopic assays showed differences in the chaotropic unfolding behaviour between SodA and SodM. However, the data from each assay, Trp fluorescence and CD spectroscopy, implied that the proteins were unfolded to different extents by the same concentration of the chaotropic agents. We therefore repeated the Trp fluorescence assays after incubating the proteins in the presence of identical guanidine concentrations as were used in the CD spectroscopy experiment. The spectra showed only a minor redshift and increase in fluorescence intensity for Fe-SodA in the presence of 4 M guanidine (Supp. Fig. 5 A), in contrast to the complete loss of α-helical structure at the same concentration (Supp. Fig. 4 A). A nearly identical redshift and increase in fluorescence intensity was observed for Fe-SodM at this same guanidine concentration (Supp. Fig. 5 B), whereas this form showed negligible loss of α-helical content in CD (Supp. Fig. 4 B). Similarly, Mn-SodA (Supp. Fig. 5 C) and Mn-SodM (Supp. Fig. 5 C) showed similar increased Trp fluorescence intensity and redshifts in the presence of 2.5 or 4 M guanidine, despite these concentrations affecting α-helical structure to very different extents (Fig. 3). We conclude that the effects of guanidine on SodFM structure is different when assessed by the proportion of α-helical content secondary structure vs polarity of the Trp environment caused by disruption of the hydrophobic central core of the folded protein.

Two spectral assays that assess the proteins’ folded state based on distinct biochemical properties, fluorescence spectroscopy to monitor the extent of burial of Trp residues within the folded structural core and CD spectroscopy to monitor the amount of helical secondary structure, were used to assess differences in the susceptibility to unfolding by chaotropic reagents of the staphylococcal SODs. The data indicated that the staphylococcal SODs differ in their susceptibility to unfolding by the chaotropic reagents, especially urea. In both SODs, the Fe-loaded form showed increased resistance to unfolding relative to the Mn-loaded form, likely a reflection of thermodynamically stronger bonds between the iron ions and the protein ligands. The MnSOD, which is present in all staphylococci, was unfolded by both chaotropic agents when loaded with either Mn or Fe, although Fe-SodA was more resistant to urea than guanidine. On the other hand, whereas the Mn-loaded form of the camSOD, which is unique to pathogenic *S. aureus* and related species, was unfolded by both agents, its Fe-loaded form showed only minimal unfolding by guanidine and was completely resistant to urea.

### Comparison of the thermal stabilities of the S. aureus SodFM enzymes

In both metal forms, the overall CD spectra of both proteins were indistinguishable (Fig. 4 A,B), consistent with their shared overall structure (5). To further investigate stability differences between the *S. aureus* SOD isozymes, we used CD spectroscopy to assess how the α-helical content of their secondary structures were affected during thermal melting from 25-95 °C by monitoring the CD signal at 222 nm (Fig. 4 C,D). MnSOD showed an obvious transition to an unfolded state at high temperature, with a near-complete loss of CD signal at 222 nm. The transition occurred in a single sharp phase in the Fe-loaded SodA (T_m_ = 70.7 ± 0.5 °C; Fig. 4 D), whereas two distinct transitions were observed for Mn-SodA (T_m_ = 57.7 ± 1.2 C and 70.0 ± 1.0 °C; Fig. 4 C). However, both forms of camSOD exhibited quite different behaviour, showing substantially less total loss of secondary structure even at high temperatures. Although a form of SodM with significantly less secondary structure was observed at high temperature, the overall loss of CD signal was less severe and the phase transition was more subtle for camSOD than for MnSOD. Determining T_m_ values for camSOD was not mathematically possible for this reason, although its thermal melt curve looked similar to that for MnSOD at temperatures <60 °C. The changes in CD signal occurred at notably lower temperature for Mn-SodM than for Fe-SodM.

**Fig. 4:**
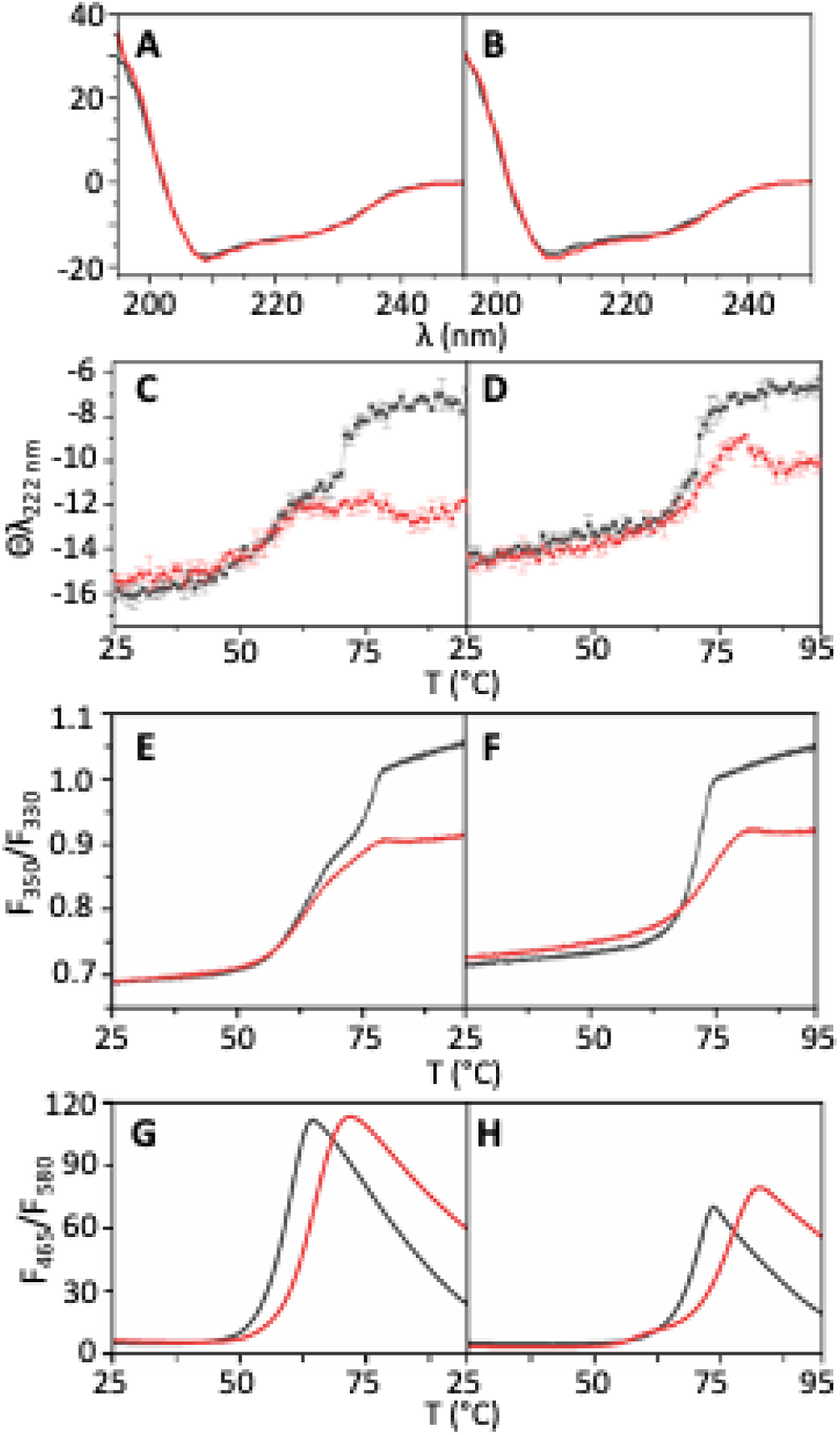
Differential thermal unfolding behaviour of the *S. aureus* SODs. Full CD spectra were obtained for the (A) manganese-loaded and (B) iron-loaded isoforms of each of the staphylococcal SOD isozymes, SodA (black) and SodM (red). CD spectra of 10 μM protein samples in 50 mM potassium phosphate buffer, pH 7.5 were recorded on a Jasco J-815 CD spectropolarimeter using 1 mm quartz cuvettes. The CD spectra of all four forms were essentially identical, consistent with very similar secondary structure content. Thermal melt curves were then obtained by measuring the CD signal at 222 nm for the (C) manganese-loaded and (D) iron-loaded isoforms of SodA (black) and SodM (red) over a temperature range from 25 to 95 °C (temperature ramp rate 1 °C min^-1^). Samples (10 μM) in 50 mM potassium phosphate buffer, pH 7.5, were analysed in technical triplicate and results presented as the average molar ellipticity (deg.cm^2^.dmol^-1^) ± standard deviation for each data point. Similar results were obtained by dynamic scanning fluorescence spectrometric analysis of (E) manganese-loaded and (F) iron-loaded isoforms of SodA (black) and SodM (red), over the same temperature range (ramp rate of 1 °C min^-1^). Samples (5 μM) in 20 mM Tris, pH 7.5, 150 mM NaCl, 5 mM EDTA buffer were analysed in technical triplicate, and the results are presented as the average fluorescence intensity ratio at 350 nm and 330 nm. Analogous results were obtained from a Sypro Orange binding assay of (G) manganese-loaded and (H) iron-loaded isoforms of SodA (black) and SodM (red), over the same temperature range (ramp rate of 1 °C min^-1^). Samples (5 μM) in 20 mM Tris, pH 7.5, 150 mM NaCl, 5 mM EDTA buffer were analysed in technical triplicate, and the results are presented as the average fluorescence intensity ratio at 350 nm and 330 nm.

To validate these data, we also performed similar thermal melt analyses using Trp fluorescence using nano differential scanning fluorimetry (DSF) to assess the unfolding of the SOD isozymes (Fig. 4 E,F). Measurement of the ratio of fluorescence at 350 (unfolded) vs. 330 nm (folded) (Fig. 1) for each SodFM isozyme at temperatures from 25-95 °C enabled us to assess the same unfolding transitions as detected in the CD, but this time using the environment of Trp residues to assess folding rather than α-helical content. The distinct biphasic unfolding of the manganese isoforms was clearly detected in an analysis of the first-derivative of the DSF data (Supp. Fig. 10), as was the extent to which the magnitude of this second thermal transition differed between SodA and SodM. The Trp fluorescence data (Fig. 4 E,F) gave similar overall thermal melting patterns as was observed by CD (Fig. 4 C,D), indicating that both assays similarly assess folding state, although interestingly the temperatures at which the transitions were observed differed more substantially in the manganese loaded isoforms, especially of SodM (Supp. Fig. 11).

Taken together, thermal unfolding studies confirmed a clear biophysical difference between the stability of MnSOD and camSOD at high temperatures, with camSOD showing increased resistance to unfolding of its secondary structure. Again, it was observed that the Fe-loaded forms of both isozymes were more resistant to unfolding than were the Mn-loaded forms.

### Overall pattern of hydrogen-deuterium exchange in the SodFM enzymes

The spectroscopic data indicated that the overall structure of the SODs was remarkably stable, resistant to unfolding at high temperatures or even in the presence of high concentrations of the chaotropic agents urea and guanidine. Crucially though, they also indicated differences in the stability of MnSOD and camSOD with respect to both thermal and chemical unfolding. To further explore the stability of the folded structure, we monitored hydrogen-deuterium exchange by mass spectrometry (HDX-MS) experiments on each of the *S. aureus* enzymes to assess the extent to which main chain amide protons in different regions of the protein are in exchange with solvent molecules, and how this differs between this closely related pair of enzymes. Samples of each isozyme, in each metal form, were exposed to deuterated solvent and samples collected over time to assess the rates of hydrogen-deuterium exchange of their alpha protons by mass spectrometry (MS). In each sample, more than 100 peptides/condition were analysed. Examination of the exchange rates in the *S. aureus* SODs identified sets of peptides with distinct rates of proton exchange: peptides that exchanged fast, exhibiting high levels of deuterium exchange even after short exposure times; those that had intermediate exchange rate, with significant exchange noted only after longer incubation with deuterated buffer; and slow exchanging regions, that exchanged little even after extended incubation. The extent of exchange within a given region was mapped onto the primary sequence of each SOD, highlighting which parts of the sequence exhibited fast vs slow rates of backbone amide exchange (Supp. Fig. 12).

Fast exchanging regions (shown in yellow, green and blue in Supp. Fig. 12) were mainly located on the surface of the enzyme and their exchange was evident after short exposure time, i.e. up to 5 min. These peptides included flexible regions of loops and turns, for example, the N-terminal 10-residue unstructured region exchanged up to 75% or more immediately after being exposed to deuterium. Intermediate exchanging regions tended to be involved in secondary structure elements, i.e. alpha helices and beta strands, in regions that were predicted to be less solvent accessible based on crystal structures, including the dimer interface. Slow or very slow exchanging peptides included short peptide(s) around the active site, particularly those regions of the sequence carrying the metal binding residues (Supp. Fig. 12).

### Comparison of hydrogen-deuterium exchange rates between MnSOD and camSOD

Comparisons were made between the two SodFM isozymes and between each metal-loaded form of each isozyme. In both cases, we observed that regions of the polypeptide that form α-helical structural elements tended to exhibit slower exchange dynamics than those in β-strands, which in turn were slower in exchange than those in loops. This observation is consistent with the α-helix structural element reducing the rate of exchange of backbone protons through its extensive hydrogen bonding (31, 32). Notably, although we found some significant differences in the rates of amide proton exchange between peptides derived from the different metal-loaded forms of either MnSOD or camSOD (Supp. Fig. 13), the overall pattern of exchange across the whole polypeptide showed negligible differences between the different metal-loaded forms (Supp. Fig. 12, comparing panels A and B or comparing panels C and D). This implies that their were detectable, but minor, differences between the metal forms in the stability of their hydrogen-bonding networks of their structural regions.

We observed more significant differences when we compared deuterium exchange rates between the two isozymes. The comparison between MnSOD and camSOD is complicated by their non-identical sequences, which result in distinct peptide fragmentation patterns. Nonetheless, given the 75% identity and 95% similarity between the two enzymes’ amino acid sequences (1), the fragmentation patterns are identical in some, and very closely matched in other structurally and functionally important regions, making it easier to compare these two SodFMs than almost any other pair. Importantly, peptides that were non-identical but similar enough to be directly compared included those carrying active site metal-binding (H27, H81, D161 and H165) and two residues known to be metal specificity-determining (G159 and L160 in the MnSOD SodA, and L159 and F160 in the camSOD SodM, collectively indicated as X_D-2_ and X_D-1_, respectively) residues (Supp. Fig. 12, panels A to D). In addition, key dimer interface residues, H31 and Y168, also locate on similar peptides in both MnSOD and camSOD, but the dimer interface loop (^125^GSG^127^) appears in different peptides. The region F112-W130 is fragmented slightly differently, containing F112-A120 and A121-W130 peptides in MnSOD compared with F112-L123 and F124-W130 peptides in camSOD (Supp. Fig. 12). This means that the GSG loop is located on a smaller peptide in camSOD than in MnSOD, which must be taken into account in comparisons between the two enzymes.

When we used this approach to compare the rates of hydrogen-deuterium exchange across the polypeptide sequence between the isozymes, we observed important differences in the exchange rates in several specific locations. The metal-coordinating residues, H27, H81, D161 and H165, are all located in regions that show slow exchange in both SODs, consistent with the active site being of limited dynamics. All three of these polypeptide regions exhibited less than 25% exchange in a 5 minute period, but over longer incubation periods, even these regions showed increasing exchange (Fig. 5). The region containing ligand H81 also contains the GGH motif, a highly conserved sequence in the SodFM1 (and mitochondrial SodFM5) subfamily consisting of mostly Mn-preferring enzymes (11). This motif is replaced by an AAQ motif in the SodFM2 subfamily, which are more frequently found to be iron-preferring enzymes (11). Notably, the region containing ligands D161 and H165 that exhibited slow exchange also contained the X_D-1_ and X_D-2_ residues that we have previously demonstrated are critical in determining the catalytic metal-preference in these enzymes and whose evolution has enabled adaptation of this critical biochemical property (1, 11). Crucially, all three of these regions showed some variation between the SodFM isozymes, indicating the core of the structure around the active site that resists exchange in camSOD was more extensive than in MnSOD. This could be seen very clearly when we mapped only the peptides that exhibited very little exchange (using a 30% cutoff) at each time point. Comparing the regions that remained below this cutoff at early (10 s) and late (24 h) timepoints in our series demonstrated how the central ‘core’ of the SodFM structure around the active site (Supp. Fig. 14, 15), which is highly hydrophobic in nature (Supp. Fig. 16) that resists solvent exchange is much larger in camSOD than in MnSOD (Fig. 6).

**Fig. 5:**
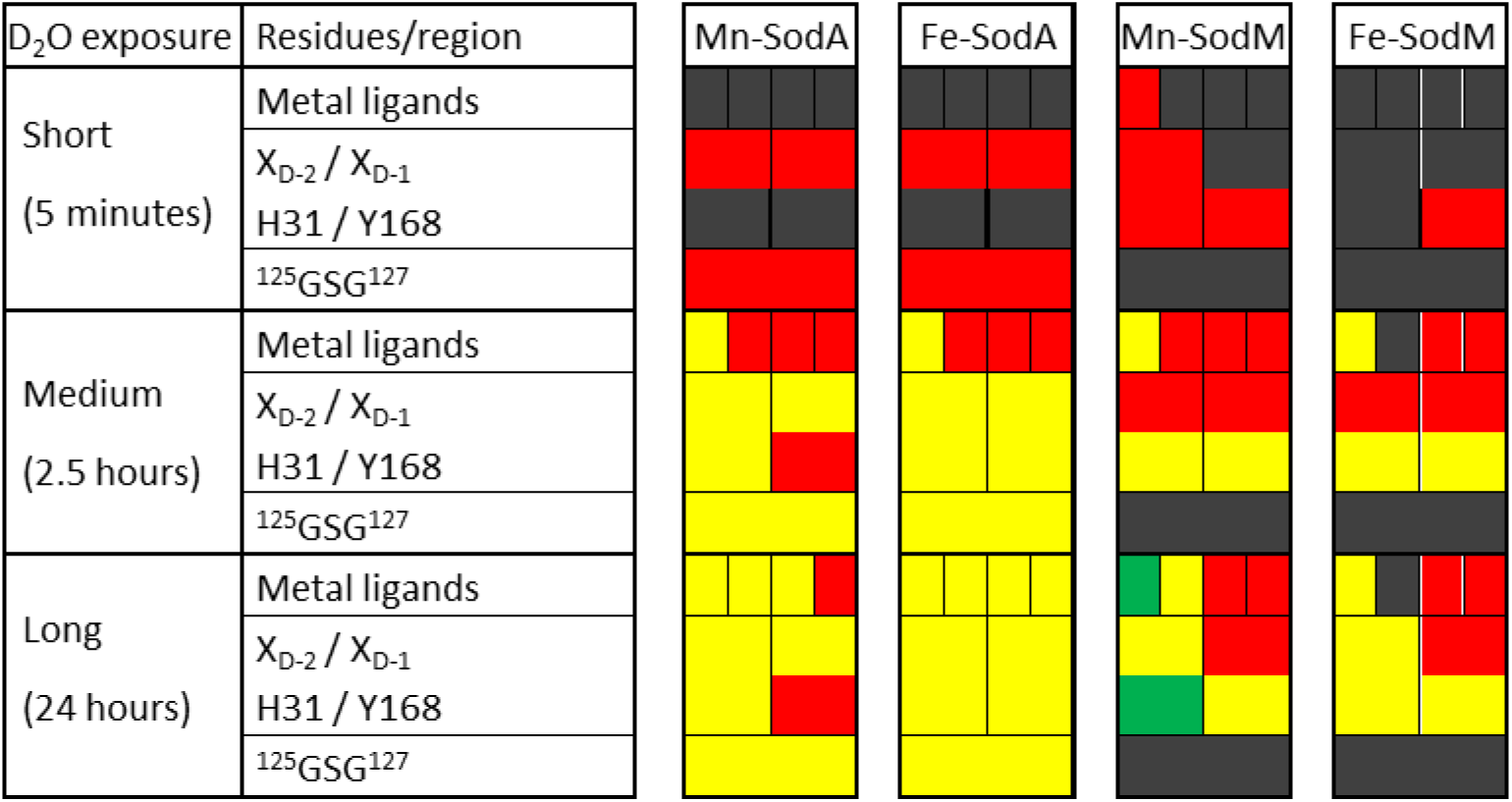
HDX-MS analysis demonstrates differences in proton exchange rates of key structural and functional residues. HDX-MS analysis of peptides derived from Mn- or Fe-loaded SodA and SodM, after short (5 min), intermediate (2.5 h) or long (24 h) exposure to deuterated solvent (see Supp. Fig. 10) was used to identify differences in main chain amide exchange rates between the two isozymes. The colour scheme shows the fraction of exchanged amide protons after a given incubation period, >12%: black, 12 – 25%: red, 25 -50%: yellow, 50 – 75%: green and 100%: blue.

**Fig. 6:**
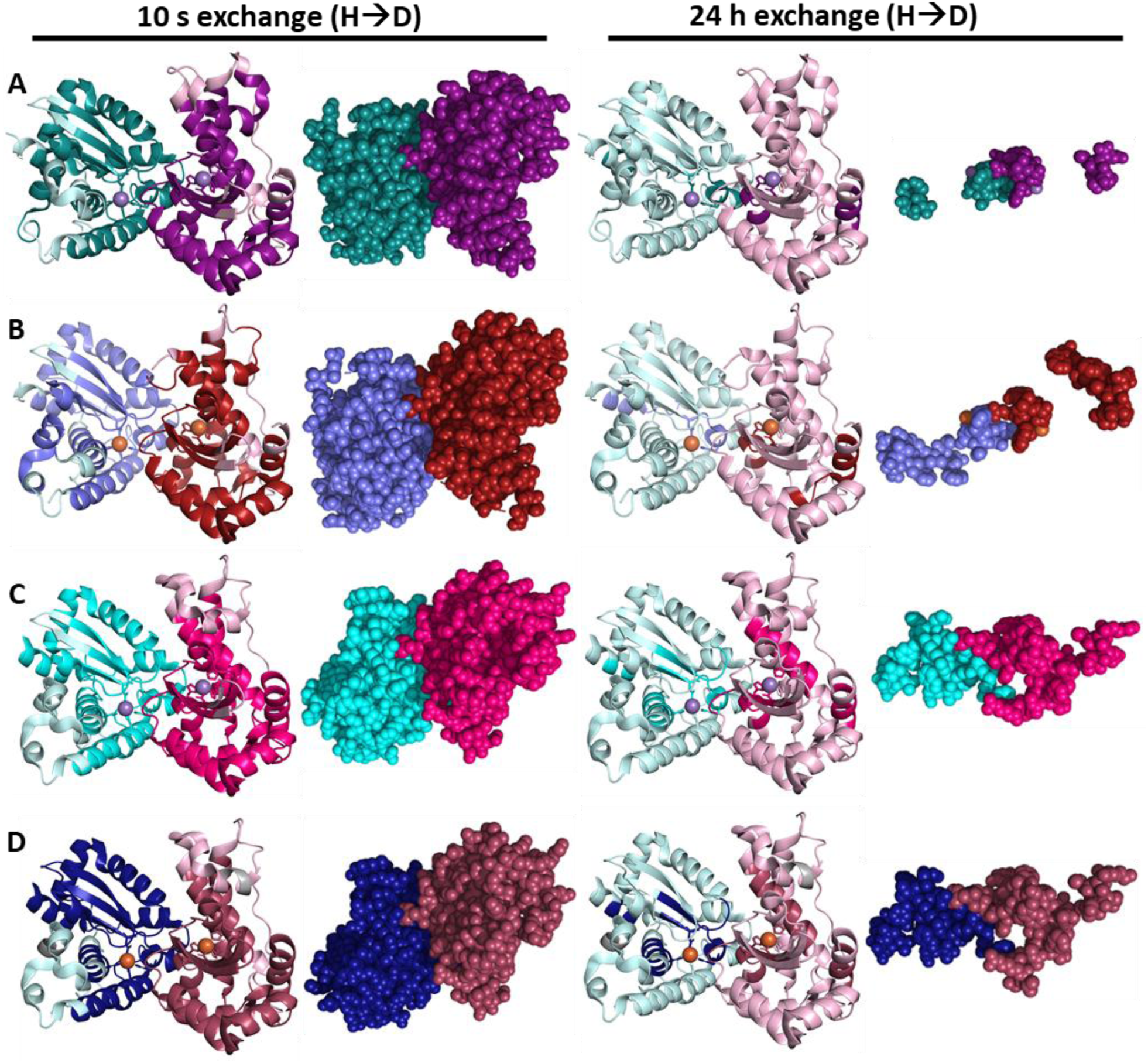
Structural distribution of regions of SOD isozymes refractory to proton exchange. Peptide regions that exhibited low overall deuteration (30% cut off) was shown on structural models to illustrate which regions of each isozyme’s structure was resistant to main-chain amide proton exchange. In each case, the illustration on the left shows the complete structural model, with regions exhibiting negligible exchange illustrated in dark colours, whereas the illustration on the right shows only those regions refractory to exchange in a space-filling representation. Given HDX provides exchange rates on the backbone amides and not the side chains, this visualization is only to present the positions of the regions with less than 30% at each time point in A: SodA Mn, B: SodA Fe, C: SodM Mn, D: SodM Fe, after very short (10 s) and very long (24 h) incubation periods in deuterated solvent.

More dramatic differences in exchange rates were observed in regions of the sequence that are involved in the dimer interface, most notably the region that included the GSG loop (^125^GSG^127^). This region showed negligible exchange in camSOD even after long incubation periods, whereas the same residues in MnSOD showed substantial exchange on the timescale of minutes to hours (Fig. 5). The GSG loop is part of a longer peptide in MnSOD than camSOD, but this difference in polypeptide fragmentation cannot explain this difference in its exchange rate. Notably, this peptide also contains two highly conserved Trp residues, W128 and W130 in a conserved WXW motif putatively important in structure and metal preference (1, 8, 33), which makes the observation that this region exchanges faster in MnSOD than in camSOD especially intriguing, especially in light of our Trp fluorescence data (Fig. 1) indicating their fluorescence is more easily affected by chaotropic unfolding. Additional residues that localise to the dimer interface include H31, which is present in the same region of the sequence and shows analogous exchange rate to the ligand H27, and Y168 (Supp. Fig. 12). The sequence region containing the latter shows broadly similar rates of exchange in both SODs, with slightly greater exchange in camSOD at multiple time points (Fig. 5). Taken together, these data suggest a potentially significant difference in the stabilities of the dimers of MnSOD compared with camSOD that warrants future investigation.

Overall, the HDX-MS data indicate that both these SodFM isozymes possess a highly stable ‘core’ region, localised around the metal-containing active site (Fig. 6), made up of regions of the polypeptide that are highly refractory to exchange of their backbone protons (Fig. 5). This suggests that this region of the structure experiences very low exposure to the solvent, even during extended incubation periods (on the order of hours) that are far in excess of the doubling time of the host bacterium (on the order of minutes). Notably, while the pattern of exchange rates in these ‘core’ regions did not show substantial variation between the different metal-loaded forms of each isozyme, they did show some significant variations between the SodFM isozymes. This difference became obvious when the regions that exhibit low exchange were mapped onto the crystal structures of the respective SodFM isozymes (Fig. 6). Despite significant sequence similarity (75% identity), a result of their recent divergence from a common ancestor, and very close structural homology between the enzymes (11), the newly neofunctionalised camSOD exhibited a greater volume of this ‘core’ region that resisted exchange of its backbone protons (Fig. 6). Whereas the exchange of this ‘core’ region that was visible after 2.5 h in MnSOD continued to undergo deuterium exchange, resulting in shrinking of the volume of the structure that remained below the 30% exchange threshold chosen for this representation, this was not the case for camSOD, which showed little change in the volume of this core between 2.5 h and 24 h of incubation in heavy water.

Taken together, our data from HDX-MS experiments demonstrate that, while differences between the isoforms of each enzyme loaded with the different metal cofactors were relatively minor, much more clear differences were observed in the rates of amide proton exchange between MnSOD and camSOD. The exchange patterns were consistent with camSOD containing a larger core region of the structure that was refractory to deuterium exchange than the core within the MnSOD structure. As these exchange rates are determined to a significant extent by the stability of hydrogen bonding networks within the protein fold (31, 32), we conclude that the overall fold of camSOD is more stable than that of MnSOD.

## Discussion

The pair of SodFM isozymes from *S. aureus* have proven to be a useful model system in which to interrogate the mechanisms by which the biochemical properties of metalloenzymes evolve (5, 11, 20). This pair of SodFMs were most likely derived from a relatively recent gene duplication event in the common ancestor of the *S. aureus* branch of the staphylococci (11). This created a second copy of the gene that encodes the highly Mn-preferring SodA (MnSOD), which is present in all staphylococci. The duplicated copy subsequently underwent evolutionary neofunctionalisation during a period of relaxed selection to yield extant SodM (20), which is present only in *S. aureus* and closely related pathobiont species (e.g. *S. argenteus* and *S. schweitzeri*) (11). Crucially, the neofunctionalisation process resulted in a gain of function, the ability of SodM to utilise either Mn or Fe as a catalytic cofactor (21). The cambialism of SodM (camSOD) is important for *S. aureus* to overcome the Mn deprivation (11, 21) it experiences during infection due to the host’s nutritional immunity response (23, 24, 34). Thus, evolution has adapted a key biochemical property of the camSOD, SodM, relative to the MnSOD, SodA, namely its catalytic metal preference. However, whether other properties of these SODs were also altered during neofunctionalisation had not yet been studied.

During our early investigations of this pair of SodFMs from *S. aureus*, we noticed that they also differed in their unfolding. While preparing their recombinant forms, we frequently obtained preparations from *E. coli* BL21 cells that contained mixtures of metals bound at the active site (21). To quantify the relative activity of their Mn- vs Fe-loaded forms, and for structural and biophysical investigations of the mechanism underlying their differing activity (5, 11), it was essential to obtain metal-pure preparations. Therefore, we developed methods for metal removal and replacement through unfolding and refolding of the recombinant proteins to obtain samples loaded exclusively with Mn (11, 21). In optimising the protocol for unfolding with chaotropic reagents, we observed that this process required more extreme conditions for camSOD than were necessary for MnSOD.

In this study, we have dissected this discrepancy, demonstrating substantial differences in the structural stability of the *S. aureus* SODs, despite their limited sequence differences. Using multiple assays of unfolding, we showed that the most recent of this SodFM pair to emerge, the camSOD SodM, exhibits increased resistance to chemical unfolding relative to its MnSOD ancestor, SodA. Notably, the resistance to chaotropic unfolding of SodM contrasts with data from other SodFMs, such as those from *Pseudoalteromonas haloplanktis* (37), which exhibited unfolding at lower concentrations of both chaotropic agent. Thermal unfolding data from CD spectroscopy, DSF and a fluorophore association assay also indicated a substantive difference in the behaviour of the *S. aureus* SODs. Consistently, SodA showed a greater extent of unfolding at elevated temperatures compared with SodM, regardless of metal-loading. In its Fe-form, SodA showed a biphasic unfolding process, with an initial transition occurring around 55 °C, followed by a further unfolding event occurring around 70 °C. This behaviour was reminiscent of that seen in CD data from other SodFMs, including the canonical Mn-preferring SodFM1 from *E. coli* (35), the cambialistic SodFM1 from *Streptococcus mutans* (36) and the Fe-preferring psychrophilic SodFM2 from *Pseudoalteromonas haloplanktis* (37). A similar pattern of thermal unfolding was observed for the closely related Mn-preferring SodFM1 from *Staphylococcus equorum* (38) and the Fe-preferring SodFM3 from the thermoacidophilic crenarchaeon *Acidilobus saccharovorans* (39). Conversely, our data showed distinct behaviour from *S. aureus* SodM, which exhibited the same initial transition at around 55 °C but lacked the subsequent unfolding event at higher temperatures. We conclude that the neofunctionalising mutations that occurred in the camSOD sequence, which imparted its flexibility to catalyse its reaction with an Fe cofactor, concomitantly altered its structural properties, creating a cambialistic SodFM that is significantly more stable than ancestral MnSOD.

HDX-MS analysis of the *S. aureus* SODs were consistent with the observed stability of the conserved SodFM structure. Protein structures are, in general, highly dynamic, leading to relatively fast rates of proton-deuterium exchange when exposed to heavy water, even for backbone protons that are buried within the hydrophobic cores of the folded protein structure (40). The exchange rates observed for backbone protons in such regions is nonetheless frequently found to be slower in metal-loaded proteins than in their corresponding apo proteins lacking metal cofactors (41–44). This occurs in regions localised to the metal-binding site, most likely through direct protection effects, but also at disparate sites, likely due to the enthalpically strong metal-ligand bonds acting as rivets that lock disparate regions of the protein together within the fold, reducing their ability to dissociate from each other through regular thermal fluctuations. Indeed, this is likely one reason for evolutionary recruitment of certain metal ions as structural cofactors in proteins. An obvious example is the employment of redox-inactive Zn(II), which forms extraordinarily high affinity bonds with protein sidechains ligands such as His or Cys. Within Zn-finger proteins, Zn(II) coordination can hold the folded protein in a rigid conformation, resistant to such conformational fluctuations, to ensure specific residues are permanently positioned correctly for interacting with its DNA target (45, 46). A reduction in conformational flexibility on binding a metal ion has even been proposed to be central to the function of bacterial metal-sensor proteins, where metal binding results in decreased conformational flexibility, excluding the conformation necessary for binding to its operator target (47, 48).

We found that both *S. aureus* SODs exhibited a central core region of the structure, constituted of regions clustered around the active site, which was refractory to exchange of its backbone protons by deuterium. Crucially, we observed subtle but significant changes in the volume of structure that resisted exchange between MnSOD and camSOD (Fig. X). As with the overall stability assays, we found that camSOD neofunctionalisation appears to have been associated with an increased strength of its hydrogen-bonded structure and/or an alteration in its conformational flexibility, resulting in camSOD exhibiting a greater volume of structure that resisted deuterium exchange.

It remains to be determined whether these properties are specific adaptations of the *S. aureus* camSOD. It’s possible that this isozyme exhibits properties that are unique among the SodFM family, perhaps a reflection of its recent emergence (20). The solvent exchange properties of the *S. aureus* MnSOD are more typical of other members of the SodFM1 subfamily, to which the *S. aureus* proteins belong, and even the entire SodFM family. Alternatively, the exchange-resistant core of camSOD might be common to, but its size highly variable among, the SodFMs. One key difference among the SodFM subfamilies is their oligomerisation, with SodFM1-2 isozymes all being homodimeric and SodFM3-5 isozymes all being tetrameric. Therefore, we predict that these different isoforms likely give rise to distinct stabilities and variation in the volume of structure that resists solvent exchange. It is interesting, in this respect, that one key region that showed variation in solvent exchange between MnSOD and camSOD lies at the dimerisation interface (residues 125-127). This raises the possibility that the dimeric interactions might be somewhat different between monomers of camSOD relative to those between monomers of MnSOD, a hypothesis that awaits future experimental testing. Nonetheless, the fact that the regions of the SodFM structure that exhibit slow exchange kinetics are noticeably altered in camSOD relative to its close paralogue MnSOD, with whom it shares 75% protein sequence identity, indicates that such properties can be evolutionarily changed through a relatively small number of mutations.

It is unclear whether the altered structural properties that we have observed in camSOD are biologically important *in vivo* and were selected during neofunctionalisation, or are merely a side-effect of the changes that adapted its metal-preference. The changes in thermal stability are evident at temperatures far beyond the optimal growth conditions of *S. aureus*, but camSOD is induced under stress conditions and thus its increased thermal/chemical stability may reflect a physiological change in enzyme stability that is functionally important under these circumstances. A previous analysis suggested that the majority (181/199) of sequence positions in camSOD were under purifying selection (20). If the increased stability is caused by a large number of mutations acting together, then it seems likely that there is some significant selection pressure responsible for the evolution of the camSOD’s physical stability. The identification of which residues that have been altered during neofunctionalisation of camSOD control its change in stability and in the solvent exchange kinetics of its structural core, and whether they overlap with the residues that control catalytic metal-preference, awaits a future mutagenesis study.

After decades of study, the structure and function of the SodFM family is relatively well understood (2). A large body of structural data is available from X-ray diffraction studies on members of this enzyme family, demonstrating that all SodFMs adopt essentially identical structures *in crystallo* (1, 2). However, little is known about the dynamic properties and structural flexibility of the SodFMs, due in large part to the presence of paramagnetic metal ions. Although the paramagnetism of a SodFM’s cofactor offers opportunities to dissect the properties of the enzyme’s active site using electron paramagnetic resonance (EPR) spectroscopy (5, 11, 49), it precludes structural determination using nuclear magnetic resonance (NMR) spectroscopy. This study is part of a wider effort, by ourselves and others, to apply biophysical assays that are unburdened by this limitation to characterise the conformational flexibility and dynamics of SodFMs in order to better understand their function at the molecular level.

## Experimental procedures

### Recombinant S. aureus SodFM production and characterisation

The protein expression and purification was as described previously (11), with modifications of the expression conditions to maximize the metal loading. Briefly, after the transformation of the target pET22 construct carrying the different SOD genes into the BL21 (DE3) Δ*sodA*Δ*sodB E. coli* strain, the cells were pre-cultured overnight in M9 media with no casamino acid mix or thiamine added, but supplemented with 1 μM of Fe. This was subcultured in M9 media supplemented with 1 μM Fe using a 1/100 (v/v) inoculation. Protein overexpression was induced by adding 100 µM IPTG, at which point the culture was supplemented with 200 µM Fe or Mn, in order to produce the Fe or Mn loaded enzymes, respectively. For cell lysis, pellets were thawed and resuspended in buffer A (20 mM Tris, pH 7.5, 100 mM NaCl) supplemented with 20 μg/mL DNase (PanReac AppliChem) and 100 μg/mL lysozyme (PanReac AppliChem) and the sample sonicated on a Soniprep 150 sonicator at 12 microns amplitude for 10 x 30 s with 1 min intervals on ice. Protein purification first used anion exchange chromatography (AEC) using a HiTrap Q FF (Cytiva) column, and a shallow gradient of 10 column volumes between buffer A and buffer B (20 mM Tris, pH 7.5, 1 M NaCl). AEC fractions were resolved on SDS-PAGE were combined and resolved by size exclusion chromatography (SEC) using a HiLoad Superdex 75 pg 16/600 (Cytiva) column in 20 mM Tris, 5 mM EDTA, pH 7.5, 150 mM NaCl buffer. To maximize the Mn saturation of the Mn loaded SodA sample for HDX-MS, an unfolding/refolding procedure using urea was carried out a prior protocol (11, 21).

### Elemental analysis by inductively coupled plasma optical emission spectrometry (ICP-OES)

Protein samples of 5 to 10 μM were digested in 32% target concentration of ultrapure nitric acid (Merk) (i.e. 1:1 v/v protein solution to 65% stock nitric acid) and diluted 5x (10x protein dilution) before measuring the metal content on a iCAP Pro ICP-OES (Thermo-Fisher Scientifics) instrument equipped with a iSC-65 autosampler, a Torch Duo (Slot) Rev 02 argon plasma torch and a Qutegra software, which was used to calculate the metal concentration. The wavelengths for Mn (257.610 nm), Fe (259.940 nm) and Zn (206.200 nm) were selected to minimize overlapping emissions. The final metal-loading values were obtained by comparing metal to protein concentration.

### Fluorescence spectroscopy

Intrinsic tryptophan fluorescence was measured on protein samples of 10 μM concentration in SEC buffer, which were incubated in buffer only or in either 8 M urea or 6 M guanidium hydrochloride (Gdn-HCl) at room temperature for 24 h. Measurements were performed at 550-600 V in 10 mm path length quartz cuvettes using a Cary Eclipse Fluorescence Spectrophotometer (Varian Inc., Agilent Technologies). Excitation and emission spectral slits were set to 5 nm width. Tryptophan residues were excited at 280 nm and the emission spectra were recorded between 280 nm and 500 nm. All measurements were corrected using their respective blank solutions (i.e. all reagents excluding the protein) on separate biological triplicate samples, and the data averaged.

### Differential scanning fluorimetry

Thermal melt curves were obtained for the Mn-loaded and Fe-loaded isoforms of both SodA (black) and SodM (red) by nano-differential scanning fluorimetry (nanoDSF) on a Prometheus NT.48 instrument (NanoTemper Technologies). Protein samples (5 μM), in 20 mM Tris, 150 mM NaCl, and 5 mM EDTA at pH 7.5, were analyzed in Prometheus standard capillaries. The temperature range was shifted from 25 °C to 95 °C at a rate of 1 °C per minute, with data acquisition according to the manufacturer’s instructions. Each sample was measured in technical triplicate, and the results are presented as the average fluorescence intensity ratio at 350 nm and 330 nm (F_350nm_/F_330nm_) for each data point.

### Circular dichroism (CD) spectroscopy

CD signals of 10 μM protein samples in 50 mM potassium phosphate buffer pH 7.5 were recorded on a Jasco J-815 circular dichroism spectropolarimeter using 1 mm quartz cuvettes (Hellma Analytics) over a temperature range from 25 to 95 °C. The temperature ramp rate was 1 °C/min, and every 1 °C the signal was measured at 222 nm. Every 5 °C, full CD spectra over 205 to 260 nm were recorded. Samples incubated o/n at 4 °C with different concentrations of Gdn-HCl (0.1 to 6 M) and urea (0.1 to 8 M) were measured for their residual CD signals over the 205 to 260 nm range. Each sample was measured in technical triplicates and results were presented as the average molar ellipticity (deg.cm^2^.dmol^-1^) ± standard deviation for each data point.

### Hydrogen-deuterium exchange mass spectrometry (HDX-MS)

HDX analyses were performed broadly according to published protocols (50). Prior to HDX reactions, non-deuterated fractions of protein served as a source of peptide lists. For this purpose, the liquid chromatography-mass spectrometry (LC-MS) analysis was carried out with all steps the same as described below for HDX runs, but D_2_O which was used for HDX was substituted by H_2_O. Peptides were identified using ProteinLynx Global Server (PLGS) Software (Waters).

HDX incubations were performed at 7 time points (10 s, 1 min, 5 min, 30 min, and 2.5 h, 12 h and 24 h) in 4 replicates with the final concentration of 5 μM, i.e. 5 μL aliquot of 50 μM protein stock was added to 45 μL of deuterated buffer (20 mM Tris pH 7.5, 150 NaCl, 5mM EDTA) at room temperature (temperature recorded separately for each measurement).

The H/D exchange reactions were quenched by moving the exchange aliquots to precooled tubes (on ice) containing 10 μL of quenching buffer (2 M glycine in 99.95% D_2_O, pD 2.4). Samples were immediately frozen in liquid nitrogen and stored at −80 °C until further measurements. Samples were thawed right before manual injection onto a nano ACQUITY UPLC system equipped with an HDX-MS Manager (Waters). Proteins were digested on a 2.1 mm × 20 mm columns with immobilized Pepsine (AffiPro) for 1.5 min at 20 °C and eluted with 0.07% formic acid in water at a flow rate of 200 μL min^-1^. Digested peptides were passed directly to the ACQUITY BEH C18 VanGuard pre-column, from which they were eluted onto the reversed-phase ACQUITY UPLC BEH C18 column (Waters) using a 10 to 35% gradient of acetonitrile in 0.01% of formic acid at a flow rate of 90 μL min^-1^ at 0.5 °C. Samples were measured on a SYNAPTG2 HDX-MS instrument (Waters) with the parameters for MS detection set as following: ESI: positive mode; capillary voltage: 3 kV; sampling cone voltage: 35 V; extraction cone voltage: 3 V; source temperature: 80 °C; desolvation temperature: 175 °C; and desolvation gas flow: 800 l/h.

Two control experiments, each in quadruplicate, were conducted to assess the minimum and maximum H/D exchange level. To obtain the minimal exchange of each peptide (𝑀𝑚ⅈ𝑛), 10 µL of a quench buffer was mixed with 45 µL of D_2_O reaction buffer prior to the addition of 5 µL of protein stocks and analyzed by LC-MS. To obtain the full deuteration (FD) exchange control (*Mmax*), the deuteration reaction was conducted in neutral pH buffer on lyophilised Sod peptides collected from immobilized pepsin column. FD sample was than processed in LC-MS system as all other samples. The control experiments were also performed in quadruplicate.

The peptide lists obtained using the non-deuterated protein samples were used to analyze the exchange data using DynamX 3.0 software (Waters). The PLGS peptide list was filtered by minimum intensity criteria of 3000 and minimal product per amino acid of 0.3. All raw files were processed and analyzed in DynamX. All MS assignments in DynamX were inspected manually. Percentage of deuteration 𝐷(%) for all peptides were calculated in Excel from exported DynamX 3.0 data, based on the formula below, which takes into account the minimal and maximal exchange of a given peptide:

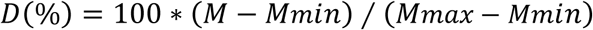

where 𝑀 is the centroid mass of a given peptide after deuterium uptake, 𝑀𝑚ⅈ𝑛 is the centroid mass of a peptide with minimal exchange and 𝑀𝑚𝑎𝑥 is the centroid mass of a peptide with a maximal exchange. Error bars for fraction exchanged represent standard deviations calculated from at least three independent experiments. Final data analysis and visualization steps were carried out using the in-house HaDeX software (51).

## Data availability

Underlying data from HDX-MS analysis is provided as a supplementary data file, and the datasets were deposited in the PRIDE repository under accession number PXD057066. Uncropped gel images are provided in the Supplementary Data. All other raw data is available from the corresponding author on request.

## Supporting information

This article contains supporting information.

## Author contributions

ME: conceptualisation, investigation, methodology, visualisation, writing - original draft, writing - review & editing; LN: investigation, methodology, visualisation, writing - review & editing; RM: investigation, writing - review & editing; LZ: investigation, writing - review & editing; MD: methodology, writing - review & editing KJW: conceptualisation, resources, supervision, project administration, funding acquisition, writing - original draft, writing - review & editing.

## Funding and additional information

ME, LN, RM and KJW were supported by a MAESTRO grant from the National Science Center (NCN), Poland (2021/42/A/NZ1/00214) awarded to KJW.

## Abbreviations

CD spectroscopy: Circular dichroism spectroscopy
EDTA: Ethylenediamine tetraacetic acid
HDX: Hydrogen-deuterium exchange
LC: Liquid chromatography
MS: Mass spectrometry
NBT: Nitroblue tetrazolium
PAGE: Polyacrylamide gel electrophoresis
SEC: Size exclusion chromatography
SOD: Superoxide dismutase
SodFM: Iron- or manganese-dependent superoxide dismutase
Trp: Tryptophan.

