## Supplementary Information for "A closely related pair of superoxide dismutase isozymes from *Staphylococcus aureus* show distinct stabilities and proton-exchange dynamics"

**Working title:** Structural stability differences between the pair of *Staphylococcus aureus* SODs

**Keywords:** Superoxide dismutase; protein stability; hydrogen-deuterium exchange; circular dichroism

**Supplementary Figures**

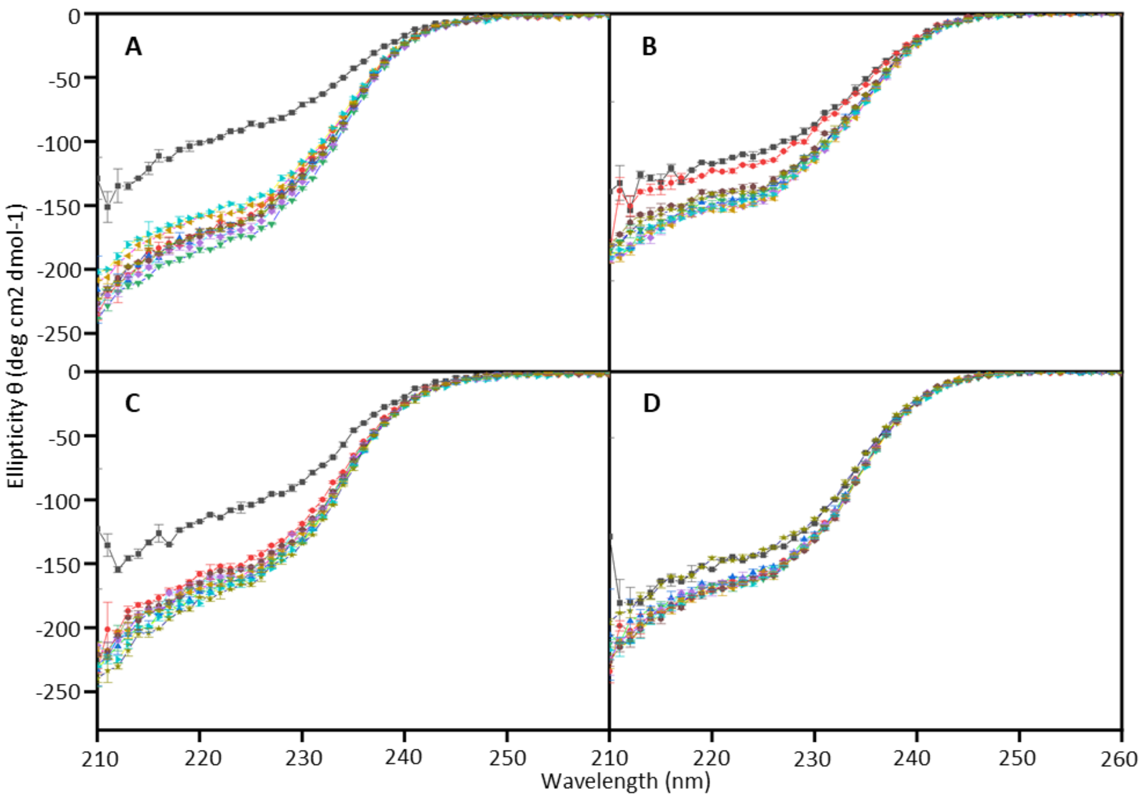


**Supplementary Figure 1: CD spectroscopy shows differential unfolding of *S. aureus* SODs in urea.** Complete CD spectra, data from which were used to produce the graphs shown in Fig. 2 and in Supp. Fig. 3. CD signals of 10 µM protein samples in 50 mM potassium phosphate buffer, pH 7.5, were recorded on a Jasco J-815 circular dichroism spectropolarimeter using 1 mm quartz cuvettes. Samples of each protein (A. Mn-SodA was determined to contain 0.983 mole equivalent Mn, 0.020 Fe, 0.001 Zn; B. Fe-SodA was determined to contain 0.990 mole equivalent Fe, 0.007 Mn, 0.000 Zn; C. Mn-SodM contained 1.070 mole equivalent Mn, 0.050 Fe, 0.00 Zn; D. Fe-SodM contained 1.090 mole equivalent Fe, 0.005 Mn, 0.000 Zn) were incubated overnight at 4 °C with different concentrations of urea (0.1 to 8 M) and then their CD spectra were over the 205 to 260 nm range

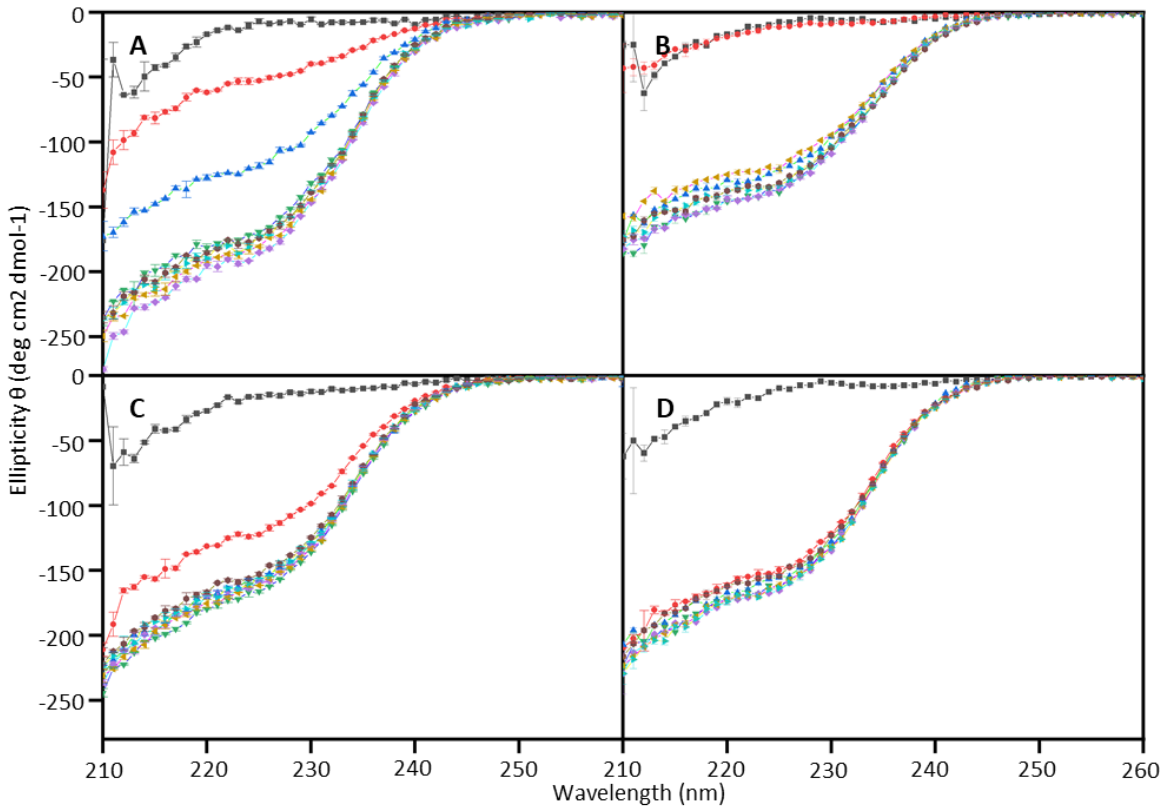


**Supplementary Figure 2: CD spectroscopy shows differential unfolding of *S. aureus* SODs in guanidine.** Complete CD spectra, data from which were used to produce the graphs shown in Fig. 3 and in Supp. Fig. 4. CD signals of 10 µM protein samples in 50 mM potassium phosphate buffer, pH 7.5, were recorded on a Jasco J-815 circular dichroism spectropolarimeter using 1 mm quartz cuvettes. Samples of each protein (A. Mn-SodA; B. Fe-SodA; C. Mn-SodM; D. Fe-SodM; metal content as described in Supp. Fig. 1) were incubated overnight at 4 °C with different concentrations of guanidine hydrochloride (0.1 to 6 M) and then their CD spectra were over the 205 to 260 nm range.

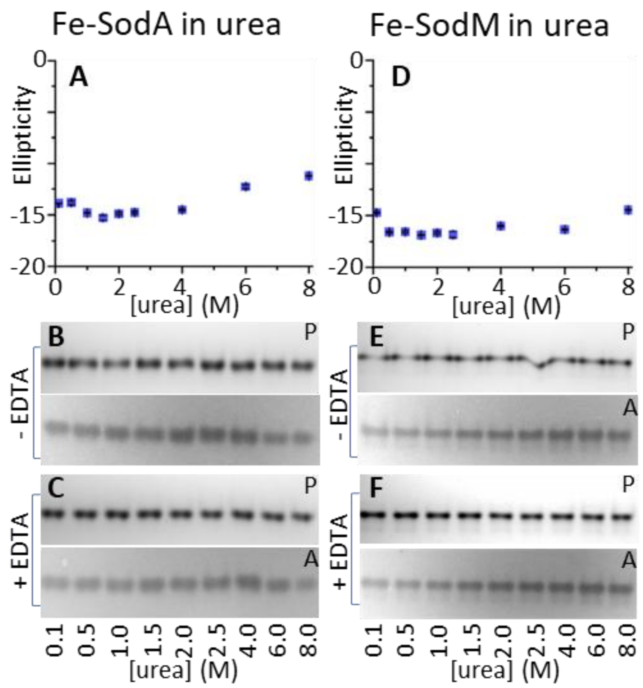


**Supplementary Figure 3: CD spectroscopy shows differential unfolding of *S. aureus* SODs in urea when loaded with iron.** Unfolding of the iron-loaded isoforms of (A-C) SodA or (D-F) SodM by the chaotropic agent urea was assessed by (A,D) measuring their CD spectra and by assessing (B,E) protein (P) stability through Coomassie staining and (C,F) enzymatic activity (A) through NBT/riboflavin staining of native PAGE gels, either before or after incubation of the samples with EDTA. CD signals of 10 µM protein samples in 50 mM potassium phosphate buffer, pH 7.5, were recorded on a Jasco J-815 circular dichroism spectropolarimeter using 1 mm quartz cuvettes Samples of each protein (metal content as in Supp. Fig. 1) were incubated overnight at 4 °C with different concentrations of urea (0.1 to 8 M) and then their CD spectra were over the 205 to 260 nm range (Supp. Fig. 1). Each sample was measured in technical triplicates and results were presented as the average molar ellipticity (deg.cm$^{2}$.dmol$^{-1}$) ± standard deviation for each data point. 50 mM EDTA (5 µL) was added to aliquots (2.5 µL) of each urea-incubated sample, incubated for 2 h, then aliquots of both control and EDTA-treated urea incubated protein samples were resolved on 10 or 12% acrylamide native PAGE, and gels stained with either Coomassie Brilliant Blue for detecting protein or with NBT/riboflavin stain to detect SOD activity. For Coomassie staining, aliquots containing 280 ng protein were loaded of all samples, whereas for activity staining, aliquots containing 698.3 ng of Fe-SodA or 41.4 ng Fe-SodM were loaded. The uncropped gel images used in this composite figure are shown in Supp. Fig. 8.

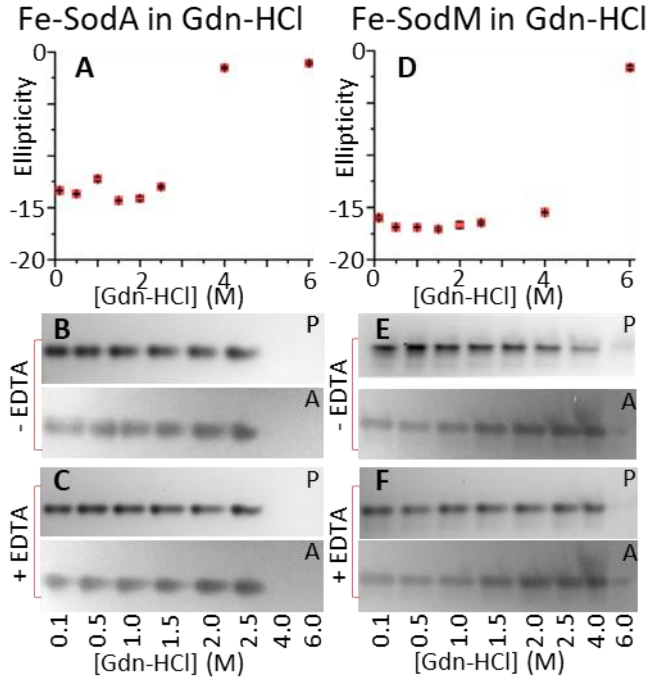


**Supplementary Figure 4: CD spectroscopy shows differential unfolding of *S. aureus* SODs in guanidine when loaded with iron.** Unfolding of the iron-loaded isoforms of (A-C) SodA or (D-F) SodM by the chaotropic agent guanidine was assessed by (A,D) measuring their CD spectra and by assessing (B,E) protein (P) stability through Coomassie staining and (C,F) enzymatic activity (A) through NBT/riboflavin staining of native PAGE gels, either before or after incubation of the samples with EDTA. CD signals of 10 µM protein samples in 50 mM potassium phosphate buffer, pH 7.5, were recorded on a Jasco J-815 circular dichroism spectropolarimeter using 1 mm quartz cuvettes. Samples of each protein (metal content as in Supp. Fig. 1) were incubated overnight at 4 °C with different concentrations of guanidine (0.1 to 6 M) and then their CD spectra were over the 205 to 260 nm range (Supp. Fig. 2). Each sample was measured in technical triplicates and results were presented as the average molar ellipticity (deg.cm$^{2}$.dmol$^{-1}$) ± standard deviation for each data point. 50 mM EDTA (5 µL) was added to aliquots (2.5 µL) of each guanidine-incubated sample, incubated for 2 h, then aliquots of both control and EDTA-treated guanidine-incubated protein samples were resolved on 10 or 12% acrylamide native PAGE, and gels stained with either Coomassie Brilliant Blue for detecting protein or with NBT/riboflavin stain to detect SOD activity. For Coomassie staining, aliquots containing 280 ng protein were loaded of all samples, whereas for activity staining, aliquots containing 698.3 ng of Fe-SodA or 41.4 ng Fe-SodM were loaded. The uncropped gel images used in this composite figure are shown in Supp. Fig. 9.

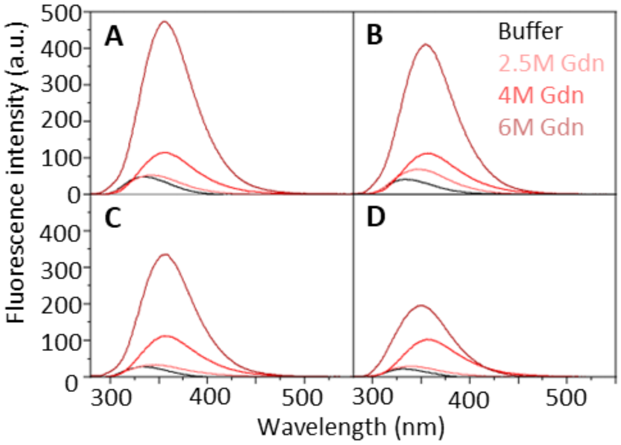


**Supplementary Figure 5: Fluorescence emission of Trp residues in *S. aureus* SODs to elaborate their differential unfolding in guanidine.** Trp fluorescence emission was assessed from samples (10 µM) of recombinant (A,B) SodA or (C,D) SodM from *S. aureus* , loaded either with (A,C) manganese or (B,D) iron. The metal loading of the samples was as described in Fig. 1. Each sample was incubated for 24 h in either buffer (20 mM Tris, 5 mM EDTA, pH 7.5, 150 mM NaCl - black), in 2.5 M (light pink), in 4 M (intermediate pink) or in 6 M guanidine (dark pink) prior to spectra being acquired. Measurements were performed at 550 V in 10 mm path length quartz cuvettes, with excitation and emission slits at 5 nm. Triplicate measurements were obtained, subtracting spectra obtained from their respective blank solutions, and averaged.

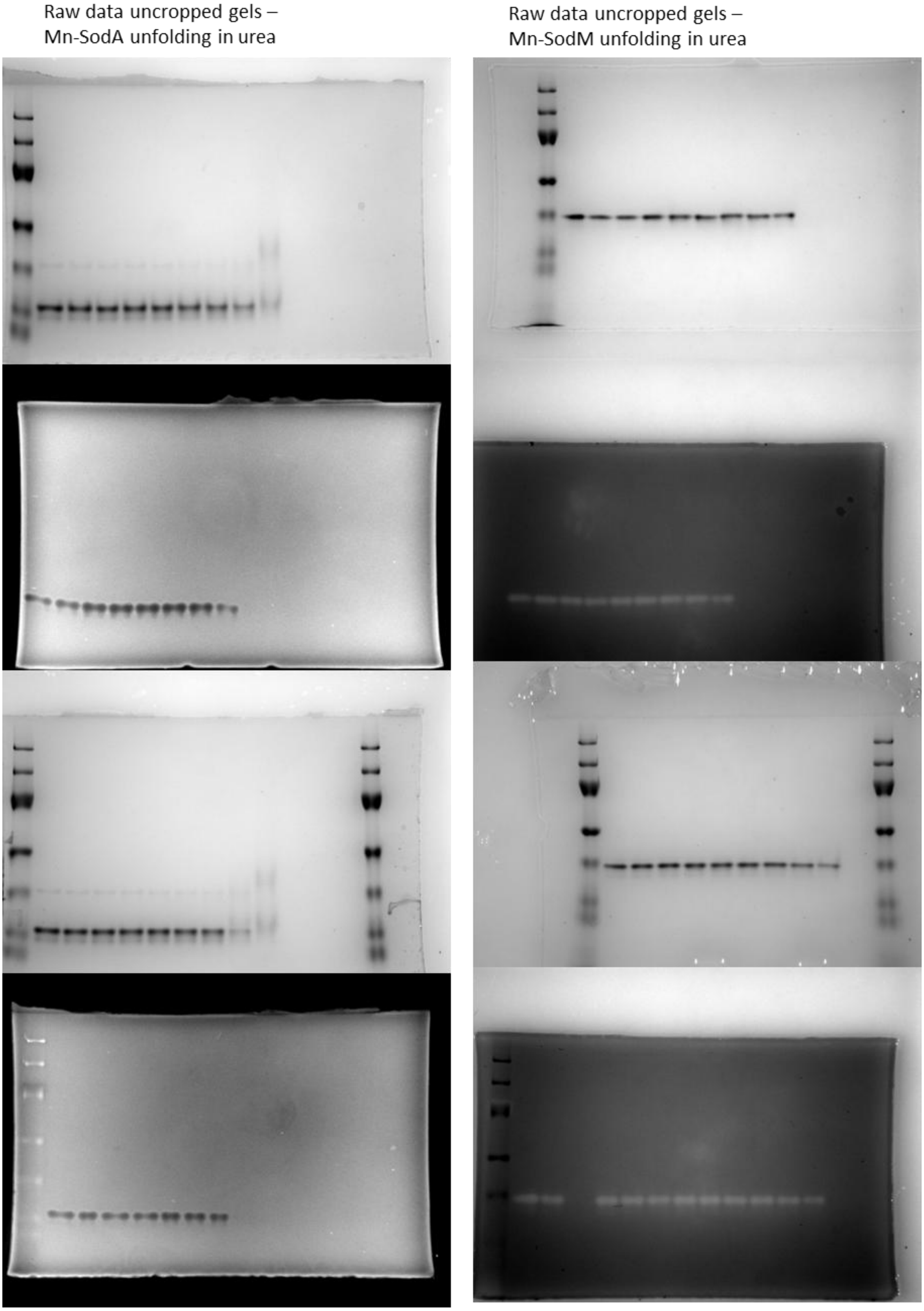


**Supplementary Figure 6: Uncropped images.** Raw data showing the unedited and uncropped gels from unfolding experiments shown in Fig. 2.

Raw data uncropped gels – Mn-SodA unfolding in Gdn-HCl

Raw data uncropped gels – Mn-SodM unfolding in Gdn-HCl

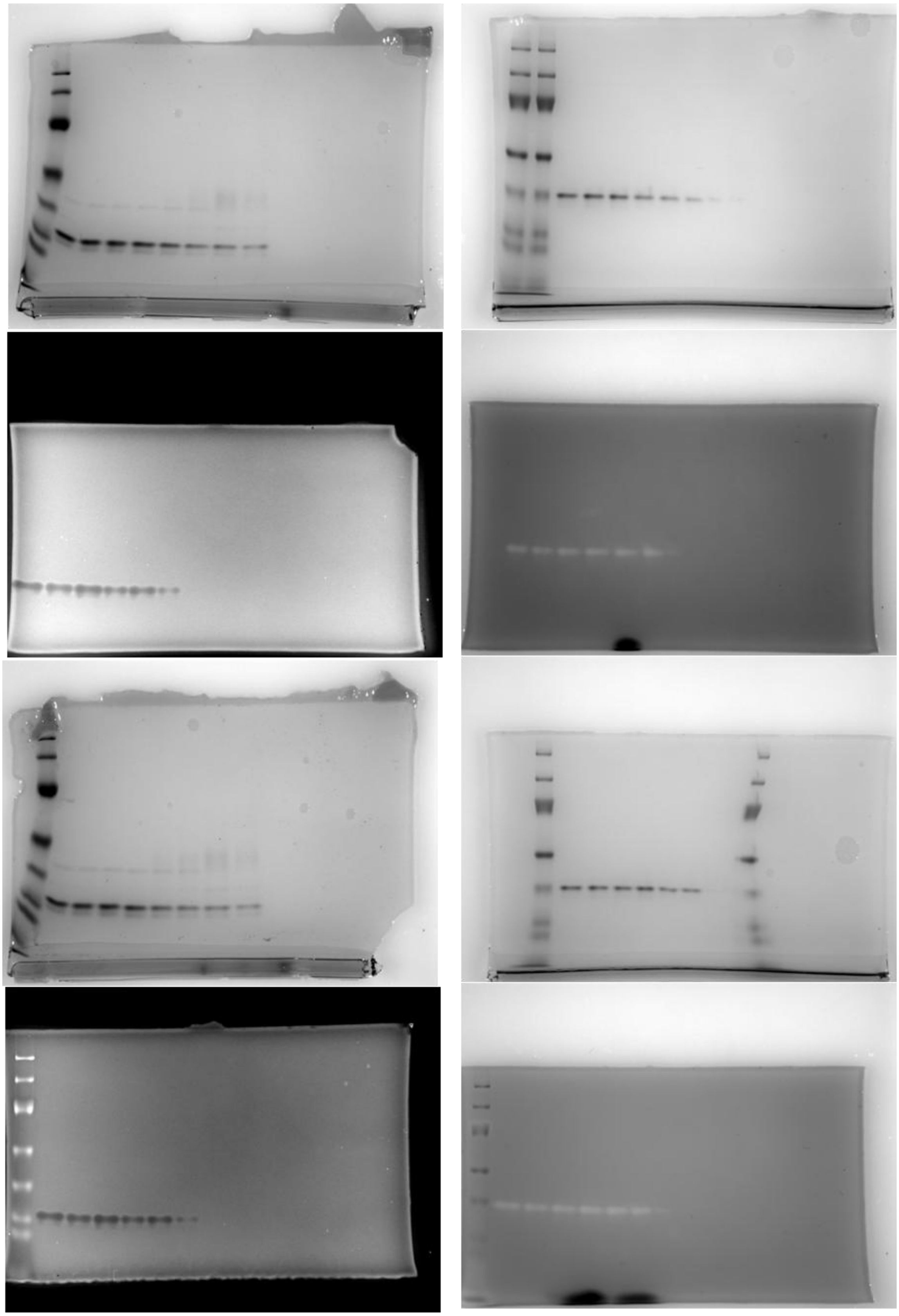

**Supplementary Figure 7: Uncropped images.** Raw data showing the unedited and uncropped gels from unfolding experiments shown in Fig. 3.

Raw data uncropped gels –
Fe-SodA unfolding in urea

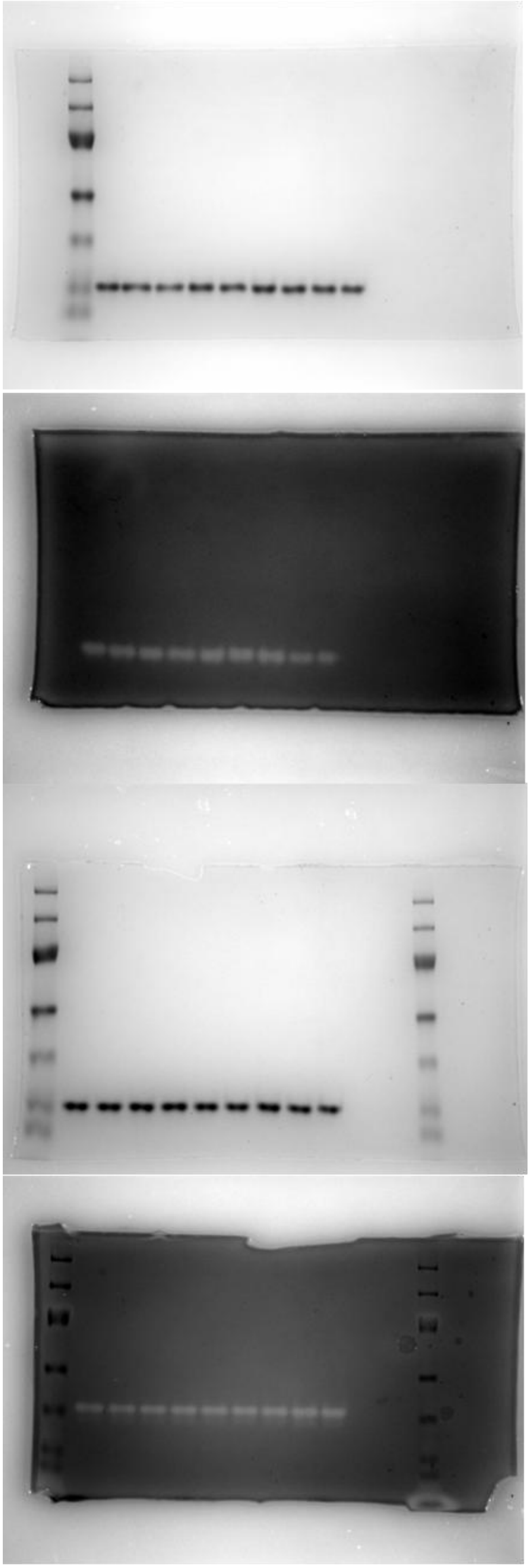

Raw data uncropped gels –
Fe-SodMunfolding in urea

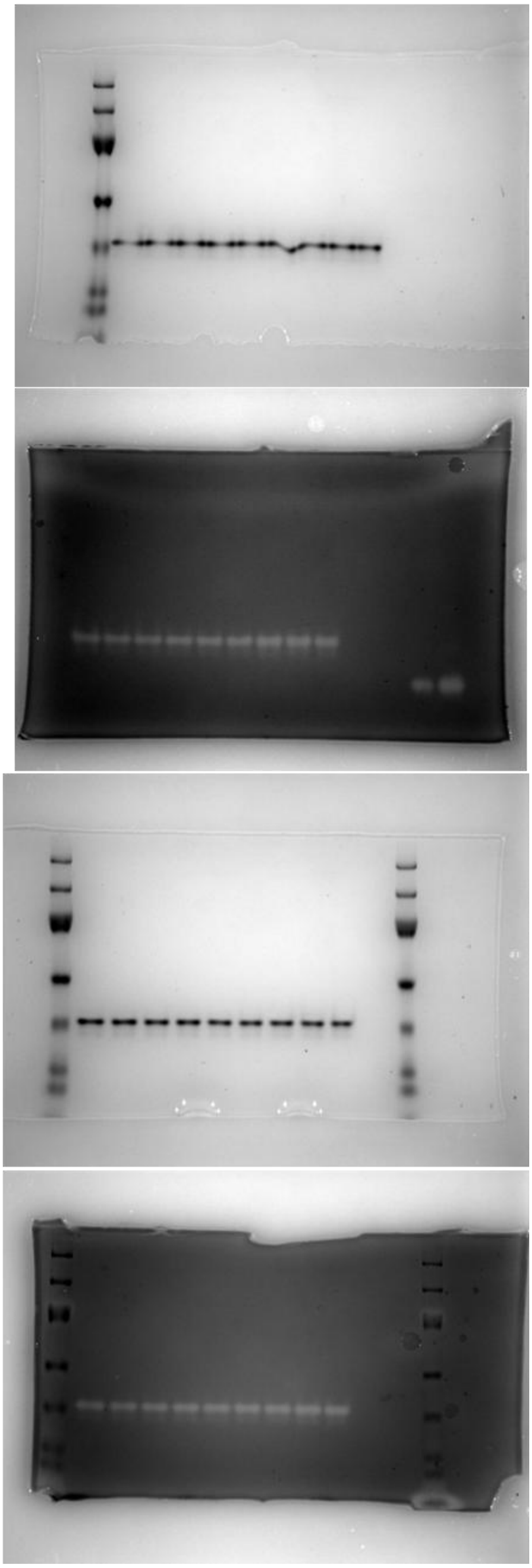

**Supplementary Figure 8: Uncropped images.** Raw data showing the unedited and uncropped gels from unfolding experiments shown in Supp. Fig. 3.

Raw data uncropped gels –
Fe-SodA unfolding in gdn-HCl

Raw data uncropped gels –
Fe-SodM unfolding in gdn-HCl

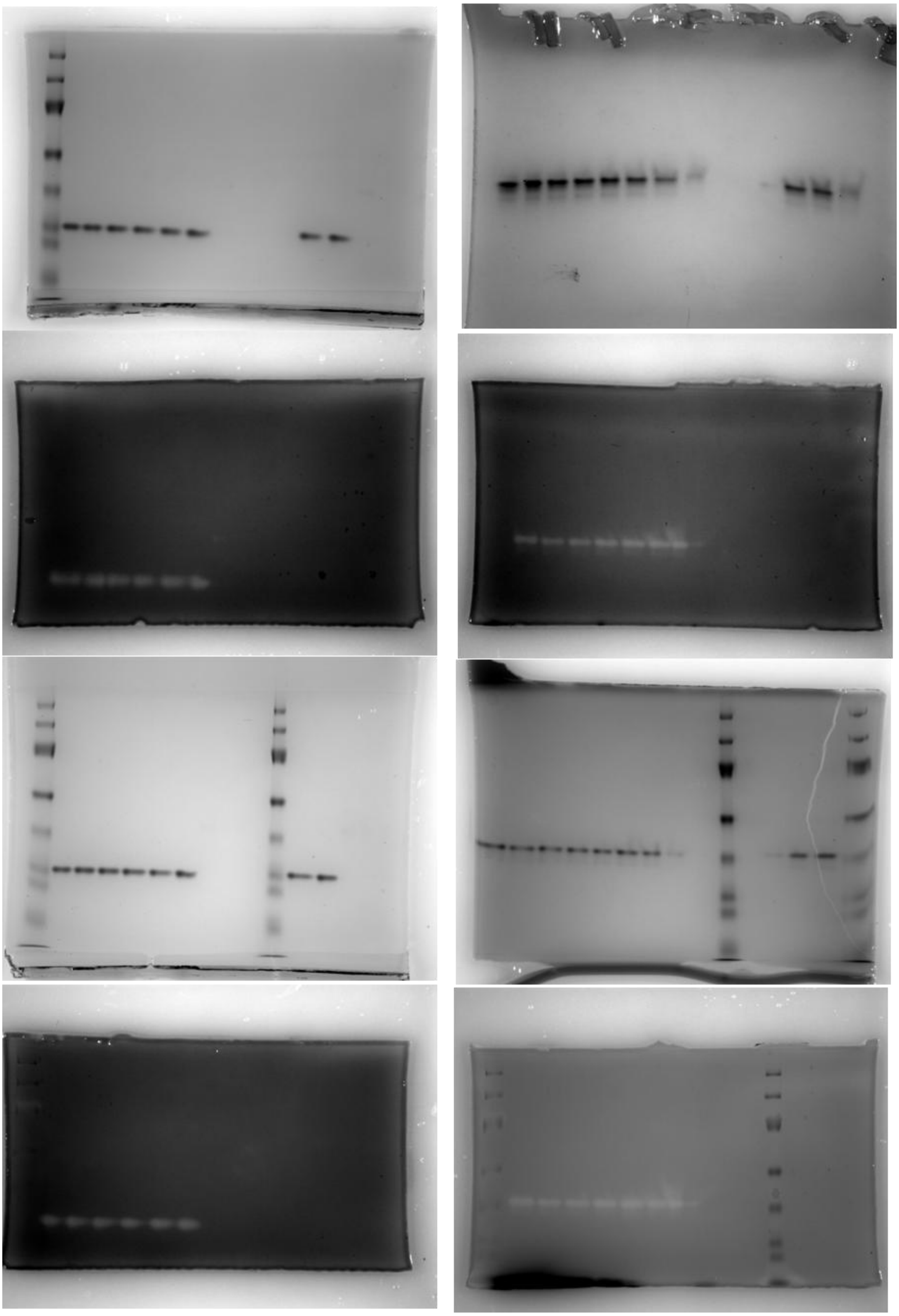

**Supplementary Figure 9: Uncropped images.** Raw data showing the unedited and uncropped gels from unfolding experiments shown in Supp. Fig. 4.

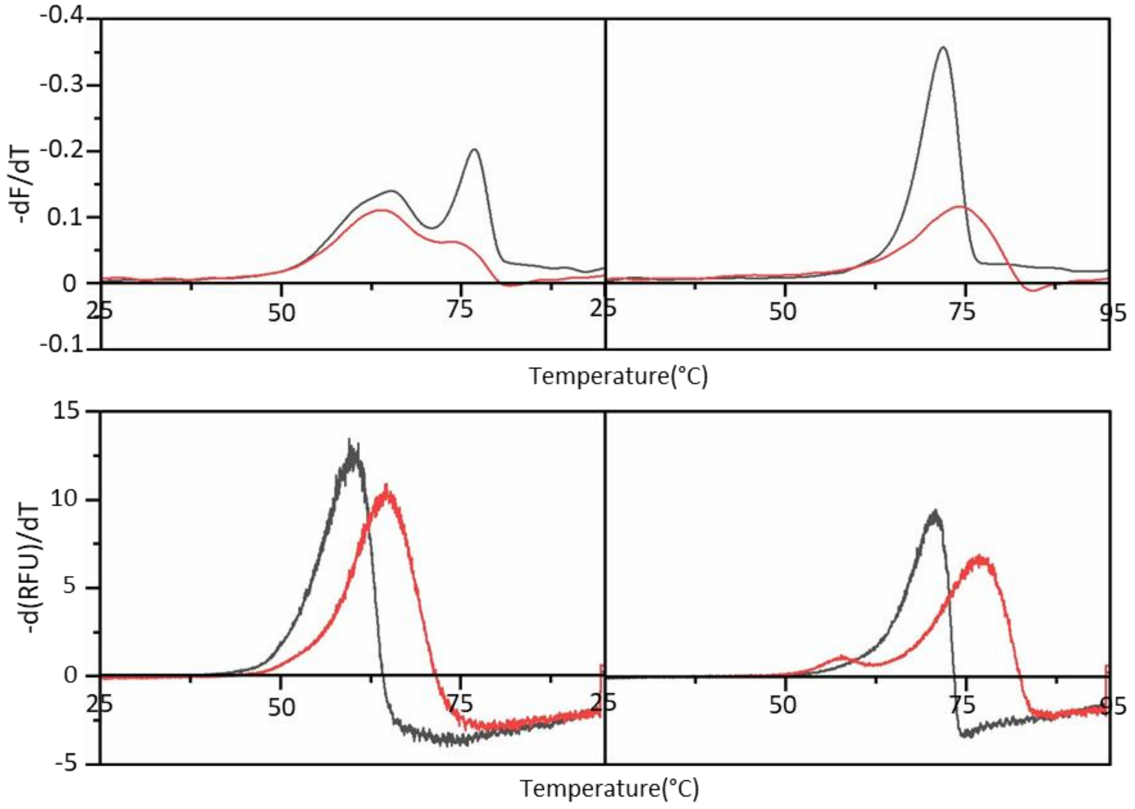


**Supplementary Figure 10: First derivative of fluorescence thermal melting curves.** First derivative curves of (upper panels) nano DSF data and (lower panels) Sypro Orange thermal assay data, illustrating the temperature-dependent structural transitions of (left) manganese loaded and (right) iron loaded isoforms of SodA (black) and SodM (red).

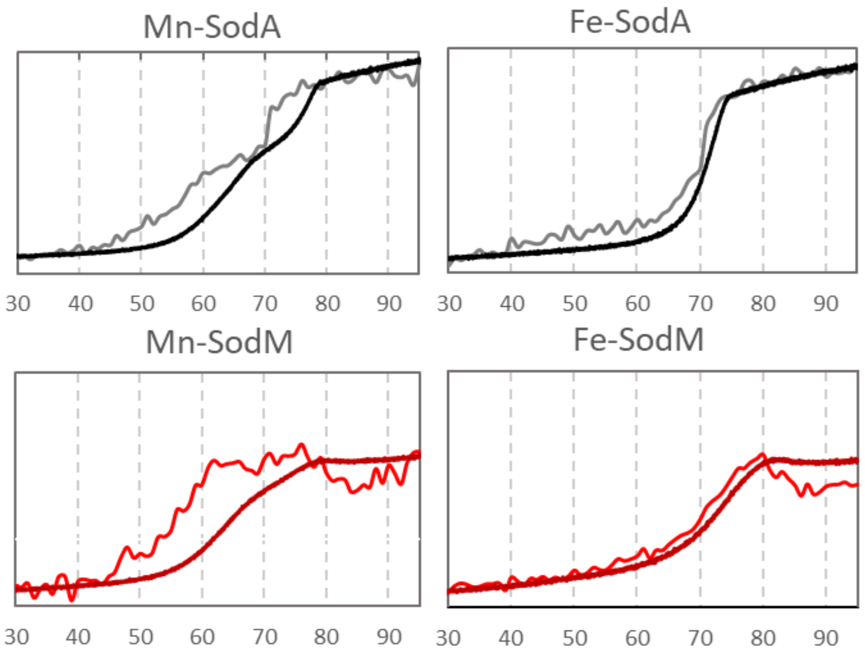


**Supplementary Figure 11: Comparison of thermal melt curves obtained from CD spectroscopy and nano DSF.** Overlaid data from CD spectroscopic analyses of protein thermal melting (thin lines), representing the deterioration of $\alpha$-helical structural content of the proteins, and nano DSF analyses of protein thermal melting (thick lines), representing the decomposition of the hydrophobic core of the structure as represented by change in Trp fluorescence. The y-axes have been normalised to allow comparison.

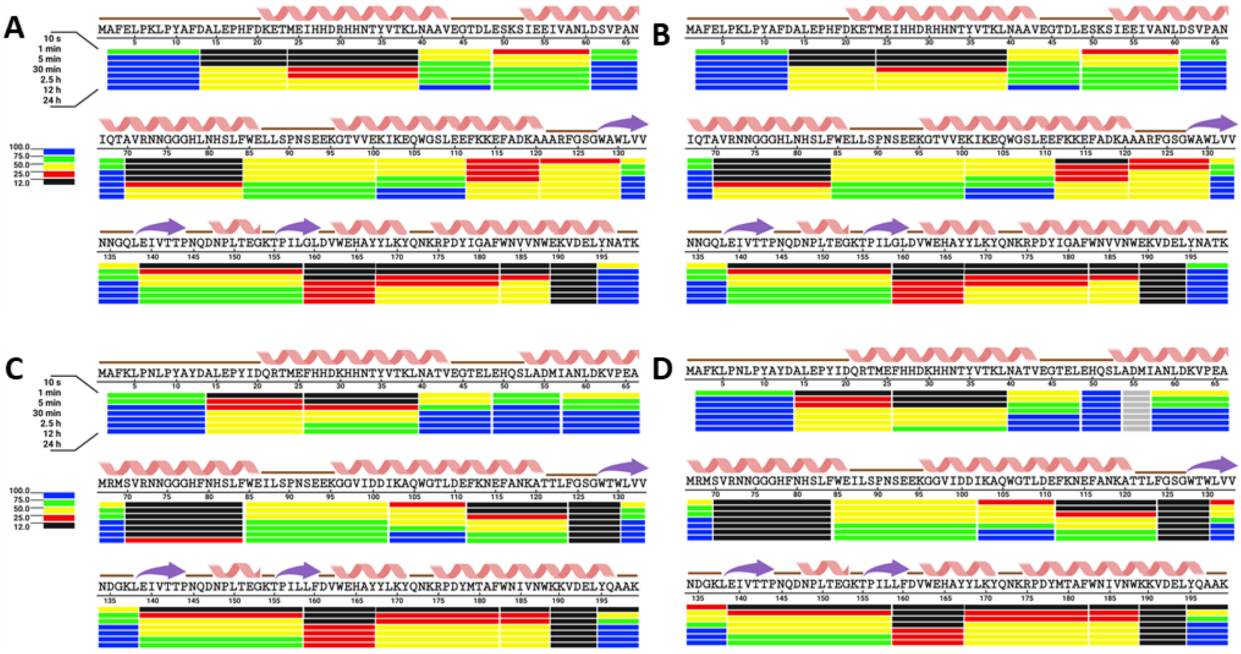


**Supplementary Figure 12: Overview of rates of hydrogen-deuterium exchange across the length of the *S. aureus* SODs polypeptides.** Overview of the pattern of rates of hydrogen-deuterium exchange observed within A) SodA Mn, B) SodA Fe, C) SodM Mn, D) SodM Fe. HDX-MS analysis of peptides was performed before (t=0 control) and after 10 s, 1 min, 5 min, 30 min, 2.5 h, 12 h, and 24 h exposure to deuterated solvent (see Supp. Fig. 10) identify differences in main chain amide exchange rates between the two isozymes. Representative peptides which enabled comparison between the SodA and SodM isozymes were selected, and their rates of exchange were depicted in BioRender. The peptides are selected to provide maximum to full coverage of the protein while retaining the shortest peptide possible in the region. Color scheme shows the fraction of exchanged amide protons after a given incubation period, >12%: black; 12 – 25%: red; 25 -50%: yellow; 50 – 75%: green; 100%: blue.

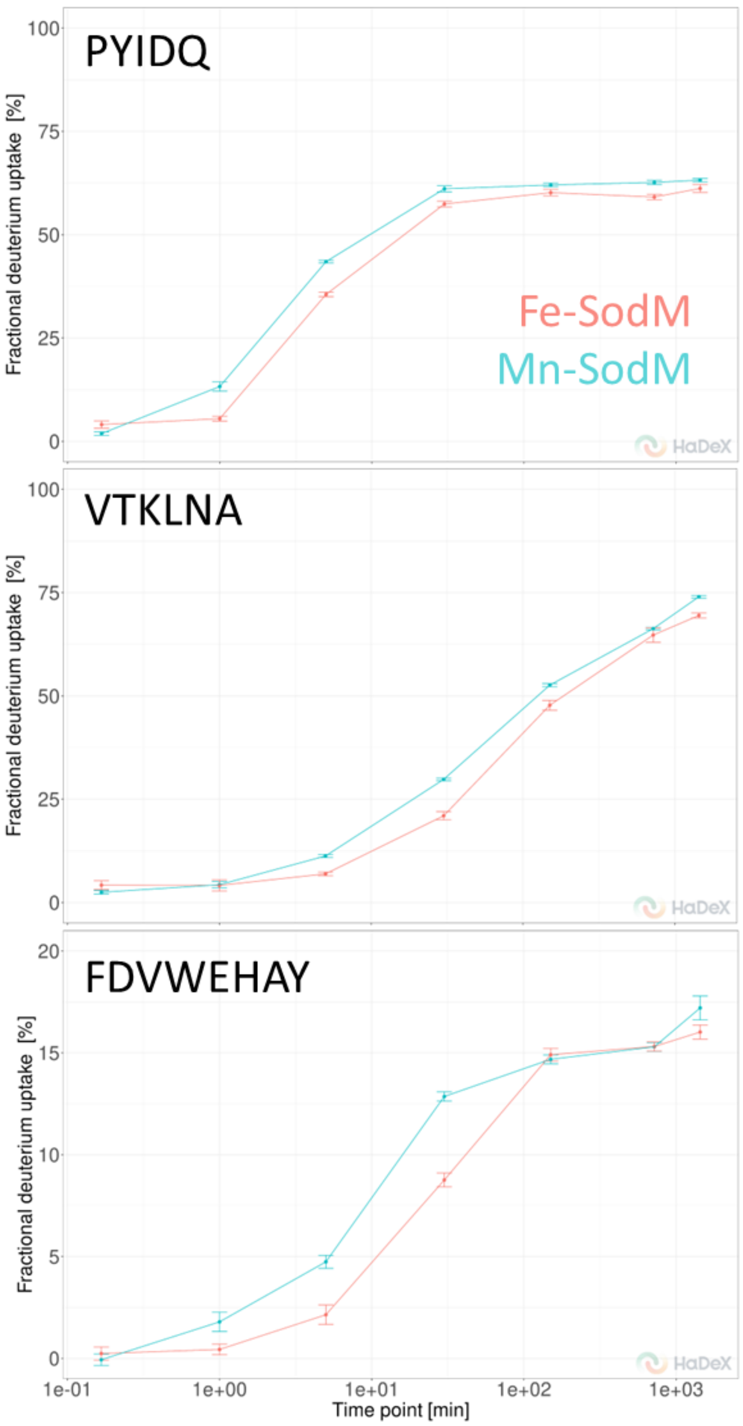


**Supplementary Figure 13: Rates of amide hydrogen-deuterium exchange in specific peptides of S. aureus SodM.** Deuterium uptake traces for three selected peptides from SodM, with data illustrating the extent of deuterium uptake in the iron-loaded (pink) and manganese-loaded (teal) form. These peptides were selected to demonstrate that some peptides did show detectable, quantitative differences in their rate of deuterium uptake between the different metal-loaded isoforms of the SodFM proteins. However, these differences were not as large as the differences observed between isozymes. The HDX-MS dataset has been deposited in the PRIDE repository under accession number PXD057066 and is publicly accessible.

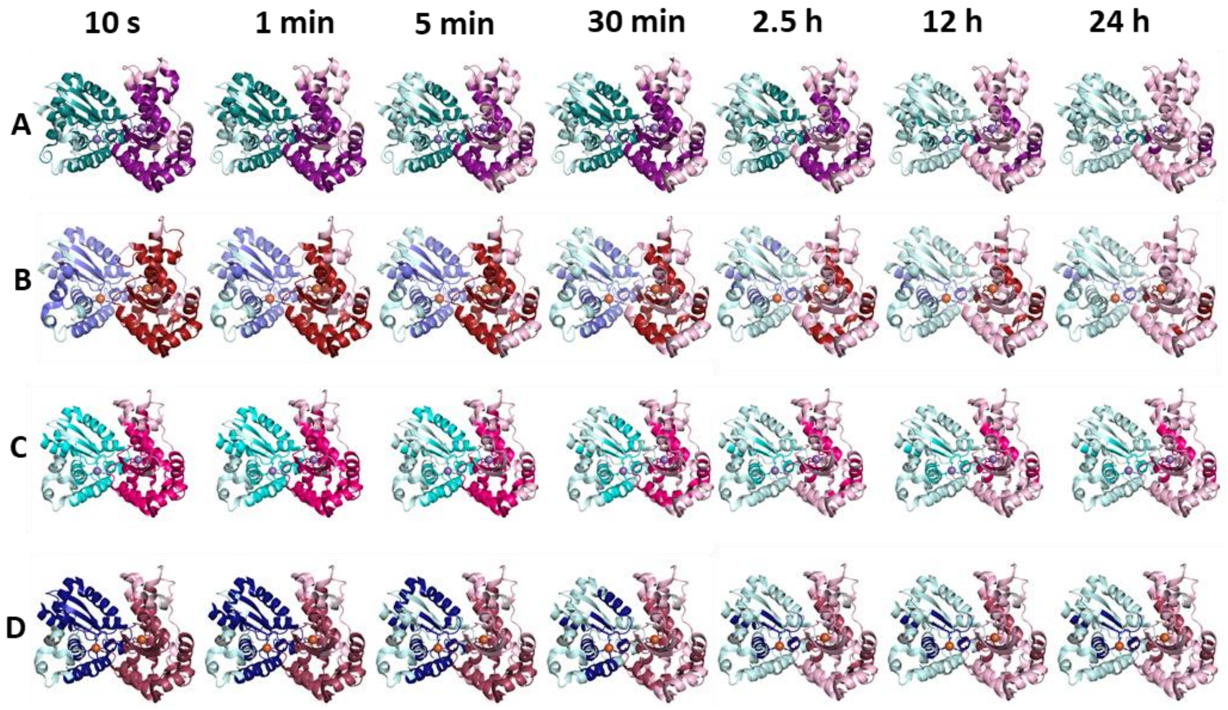


**Supplementary Figure 14: Overview of rates of hydrogen-deuterium exchange across the length of the *S. aureus* SODs polypeptides.** Structural illustration, demonstrating regions of the SodFM structures of A: Mn-SodA, B: Fe-SodA, C: Mn-SodM, D: Fe-SodM that exhibited low rates of deuterium exchange of their main-chain amide hydrogen atoms in HDX-MS analyses. Peptide regions that exhibited low overall deuteration (30% cut off) at each time was shown (dark coloured ribbons) on the overall structural models (light coloured ribbons) to illustrate which regions of each isozyme's structure was resistant to main-chain amide proton exchange. The values to represent in the structure are obtained for each residue based on the uptakes from overlapping peptides using the weighted approach in DynamX 3.0).

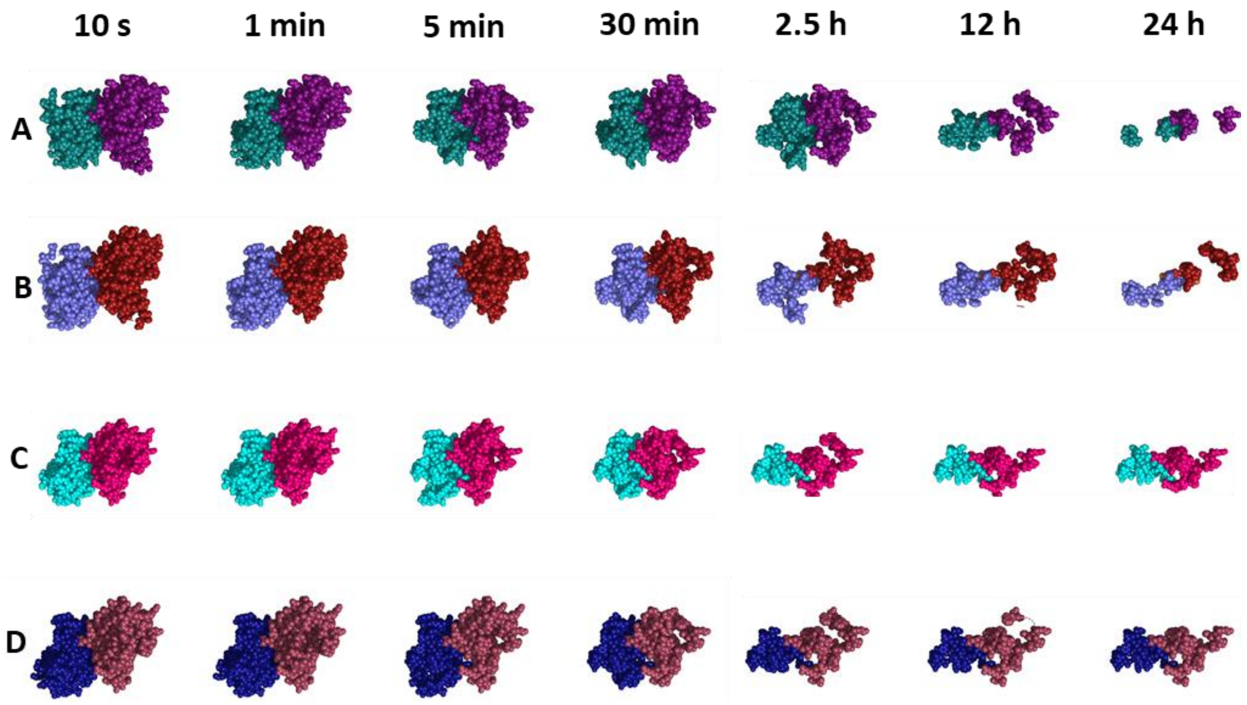


**Supplementary Figure 15: Overview of rates of hydrogen-deuterium exchange across the length of the *S. aureus* SODs polypeptides.** Alternative structural illustration, demonstrating regions of the SodFM structures of A: Mn-SodA, B: Fe-SodA, C: Mn-SodM, D: Fe-SodM that exhibited low rates of deuterium exchange of their main-chain amide hydrogen atoms in HDX-MS analyses. Peptide regions that exhibited low overall deuteration (30% cut off) at each time was shown in space-filling representation to illustrate the volume of the regions of each isozyme's structure that was resistant to main-chain amide proton exchange. The values to represent in the structure are obtained for each residue based on the uptakes from overlapping peptides using the weighted approach in DynamX 3.0).

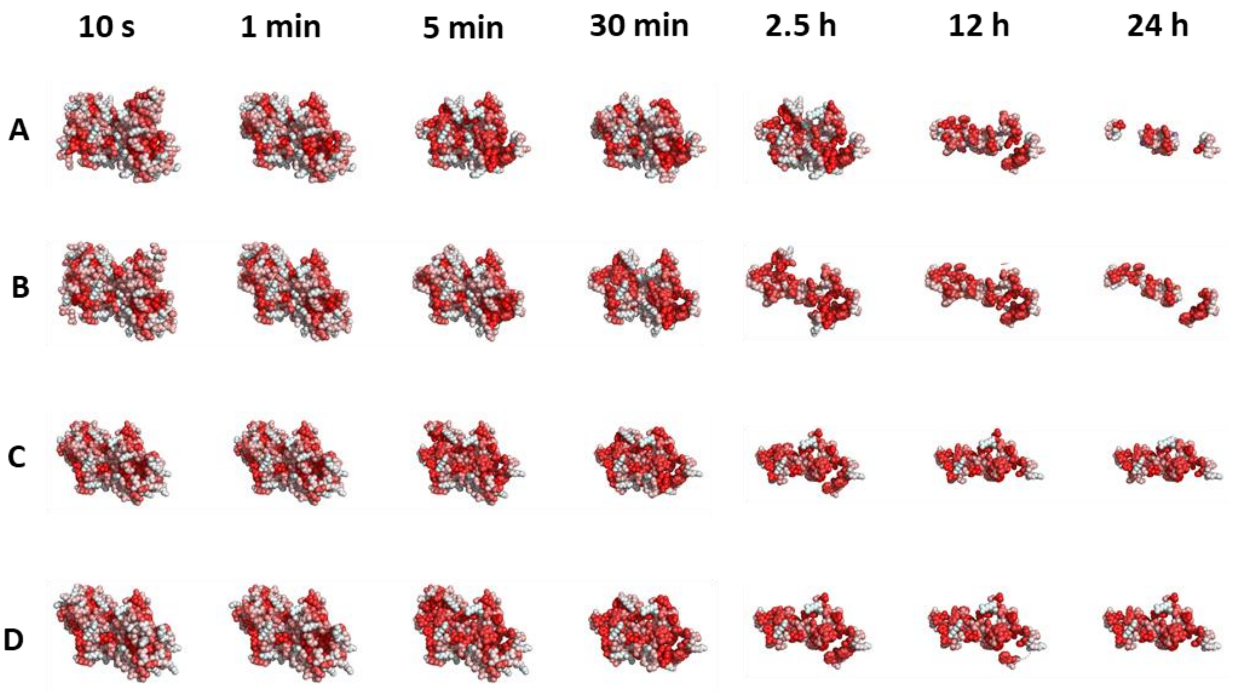


**Supplementary Figure 16: Overview of rates of hydrogen-deuterium exchange across the length of the S. aureus SODs polypeptides.** Alternative structural illustration, demonstrating regions of the SodFM structures of A: Mn-SodA, B: Fe-SodA, C: Mn-SodM, D: Fe-SodM that exhibited low rates of deuterium exchange of their main-chain amide hydrogen atoms in HDX-MS analyses. Peptide regions that exhibited low overall deuteration (30% cut off) at each time was shown in space-filling representation, coloured according to hydrophobicity in Pymol, as an alternative representation of the volume of the regions of each isozyme's structure that was resistant to main-chain amide proton exchange. The values to represent in the structure are obtained for each residue based on the uptakes from overlapping peptides using the weighted approach in DynamX 3.0).
